# MODULATION OF COLLAGEN-BINDING INTEGRINS AFFECTS FIBROBLAST ACTIVATION AND INHIBITS FIBROSIS

**DOI:** 10.1101/2025.05.14.653428

**Authors:** Stefan Hamelmann, Fatemeh Derakhshandeh, Tom Wagner, Stephan Uebel, Barbara Steigenberger, Claire Basquin, Sebastian Fabritz, Jutta Schroeder-Braunstein, Guido Wabnitz, Inaam A. Nakchbandi

**Affiliations:** Institute of Immunology, Heidelberg University, 69120 Heidelberg, Germany; Max-Planck Institute for Biochemistry, 82152 Martinsried, Germany; Max-Planck Institute for Medical Research, 69120 Heidelberg, Germany

**Author notes:** One sentence summary: collagen-integrins control fibroblast activation and can be pharmacologically manipulated to inhibit fibrosis. Shared first authorship (Stefan Hamelmann, Fatemeh Derakhshandeh). Funding sources: German Research Council (DFG) (NA-400/5-1; NA-400/5-2; NA-400/7; NA-400/9); Max-Planck Society (M.KF.A.BIOC0001/K440). Correspondence:* Inaam Nakchbandi, MD, FACP, Institute of Immunology, University of Heidelberg, Im Neuenheimer Feld 305, 2. OG, 69120 Heidelberg, Germany.

**Keywords:** Extracellular matrix, collagen, fibroblast, activation, lung fibrosis, liver fibrosis, integrin α10β1, integrin α11β1

## Abstract

The extracellular matrix contributes to the progression of several diseases, sometimes by disrupting organ function such as in lung and liver fibrosis. Because integrin receptors mediate cell-matrix interactions, we conditionally deleted β1 integrin in murine hepatocytes *in vivo*. Increased TGF-β and matrix deposition ensued. Application of a cyclic peptide (GLQGE) that binds to both α10β1 and α11β1 diminished fibrosis in two murine models. In liver fibrosis, TGF-β production was reduced. In lung fibrosis, however, the effect was exclusively due to suppressing fibroblast activation and hence collagen production without TGF-β involvement.

In summary, integrin manipulation successfully changed cell behavior, with effects differing depending on the cell type. Importantly, it is possible to directly suppress fibroblast activation and consequently diminish matrix production independent of disease pathogenesis. This underscores the importance of matrix composition in modifying the behavior of embedded cells and hence disease severity.

**Key findings:**

- Loss of α11β1 integrin-mediated signal in hepatocytes increases TGF-β and leads to fibrosis
- Engaging α10β1 and/or α11β1 with a cyclic peptide (GLQGE) inhibits fibroblast activation directly and, consequently matrix production and progression of fibrosis

## Introduction

The extracellular matrix accumulates to various degrees in many disease entities. It is usually viewed as an irrelevant byproduct of the disease except when its amount is large enough or its distribution erratic enough to disrupt organ function ^1,2^. One example is the liver, where accrual of matrix interferes with the filtering function of the liver and leads to insufficient exchange between the bloodstream and the hepatocytes ^3^. Another example is the lung, where matrix buildup diminishes O_2_/CO_2_ exchange between the air along the alveolar wall and the blood in the capillaries of the lungs ^4^. Despite major advances, no breakthrough in the treatment of fibrosis could be achieved.

The extracellular matrix is comprised of a large number of molecules, that not only provide the scaffold giving the organs their characteristic forms, but also interact with the cells they surround after being produced by them. Their importance can be deduced from the fact that universal knockouts sometimes lead to embryonic death ^5^. Fibronectin is such a molecule that not only retains growth factor TGF-β in the matrix and regulates its activation, but also affects cell survival, proliferation, migration, differentiation and activity ^6-11^. Two functions of fibronectin highlight the importance of matrix molecules. On one hand, interfering with fibronectin fibril formation prevents collagen accumulation *in vitro* ^12^. Therefore, a molecule that hinders fibril formation called pUR4 is able to diminish experimental liver fibrosis ^13^. On the other hand, conditional deletion of fibronectin postnatally in the liver results in an increase in TGF-β release, a major profibrotic cytokine. This TGF-β then stimulates the cells that produce the matrix in the liver, the so-called hepatic stellate cells or myofibroblasts leading to increased collagen type I production and fibrosis progression ^14^. While at first glance these two findings may seem contradictive, they suggest that the composition of the matrix is critical for how the various cells respond to it.

A major class of matrix receptors that mediate the interactions between cells and their surrounding are the integrins. These consist of pairs of an α and a β subunit that together bind to the matrix and affect an intracellular response. There are at least 18 α subunits and 8 β subunits. Of these, the β1 subunit plays a central role, because it can pair with 12 α subunits ^15^. While fibronectin binds to several integrin pair combinations, the classical fibronectin receptor, α5β1 integrin, contains the β1 subunit^16^. Collagen type I, the major matrix protein increased in fibrosis, can only bind to integrin pairs that contain the β1 subunit, namely α1β1, α2β1, α10β1 and α11β1 integrins ^15^. Indeed, the importance of the β1 subunit in the liver was highlighted in work showing its requirement for regeneration ^17^. We therefore hypothesized, that if the matrix modulates the severity of fibrosis, β1 integrin might be involved.

In this work, we manipulated β1 integrin and took advantage of the identified processes to suppress fibroblast activation and matrix production pharmacologically and diminish liver and lung fibrosis.

## Results

### Deletion of β1 integrin enhances matrix production and leads to fibrosis

β1 integrin pairs with four different α subunits to attach to collagen and five subunits to bind fibronectin ^18^. We hypothesized that recognition of the matrix by β1 integrin might affect matrix production. Therefore, we deleted β1 integrin in hepatocytes. This was achieved by using the cre recombinase gene under the control of the albumin promoter in mice homozygous for a floxed β1 integrin gene. The conditional knockout (cKO) mice (Alb-cKO: Alb-cre_β1^floxed/floxed^) were compared to littermate controls lacking cre (CT)^9,10^. Since albumin is only expressed in hepatocytes, deletion of β1 integrin was hepatocyte-specific ^14^. Deletion was confirmed in the liver by western blotting and in isolated hepatocytes both by western blotting and flow cytometry (Figure 1A-B).

**Figure 1.**
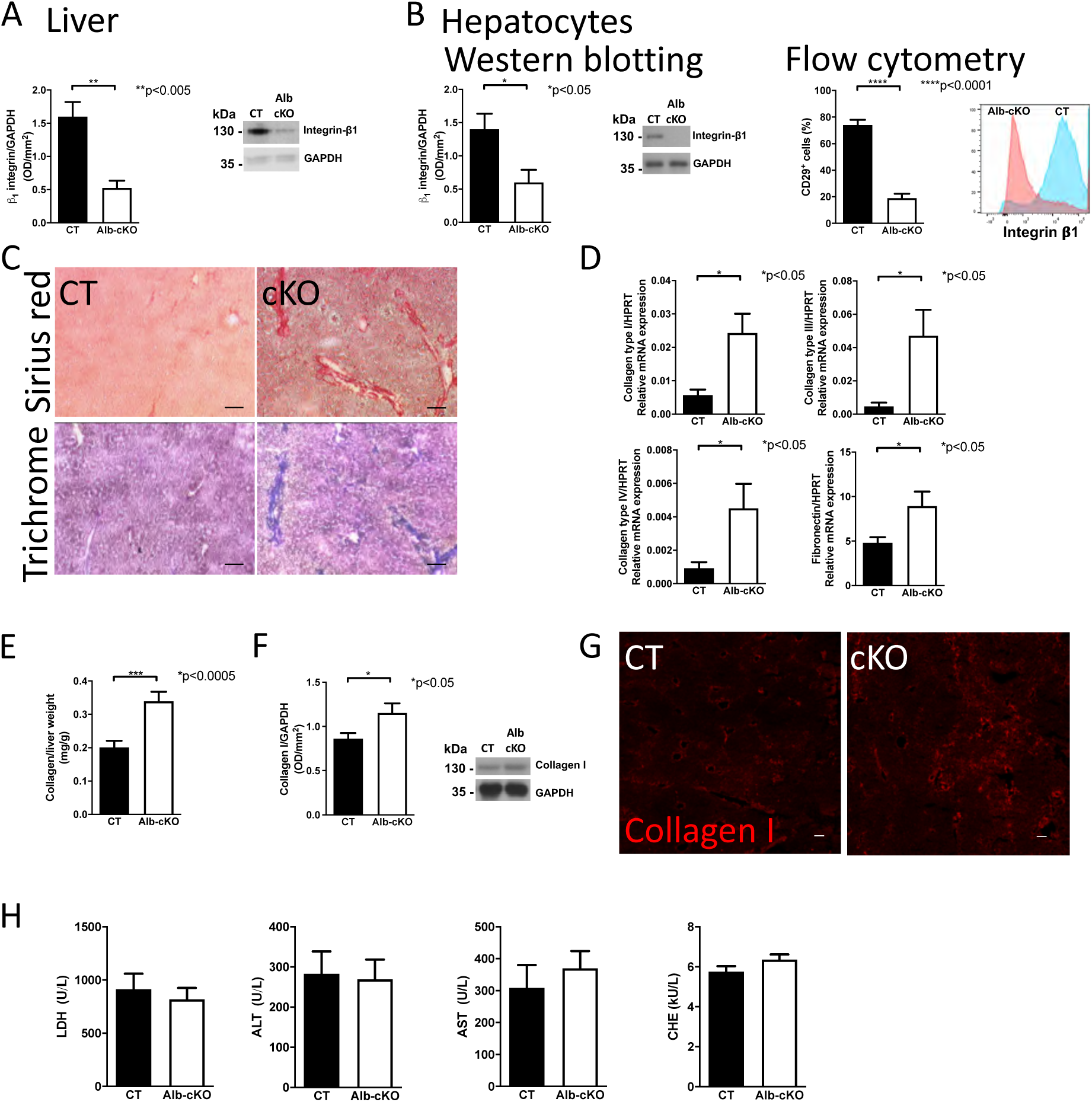
Deletion of β1 integrin in hepatocytes results in fibrosis. (A) β1 integrin is diminished in the liver of conditional knockout (cKO) animals carrying the albumin promoter to drive cre expression in mice homozygous for floxed β1 integrin (Alb-cKO) compared to control mice (CT) by western blotting. N=4 CT and 4 Alb-cKO replicates. **p<0.005. (B) Western blot analysis shows successful depletion of β1 integrin in hepatocytes isolated from Alb-cKO animals. Flow cytometry confirms a decrease in the percentage of cells expressing β1 integrin in Alb-cKO hepatocytes. The number of replicates for the groups is shown in the order presented in the graphs: N=12/10 for western blot and 6/6 for flow cytometry. *p<0.05, ****p<0.0001. (C) Sirius-red and trichrome staining suggest an increase in extracellular matrix in Alb-cKO liver sections. Bars represent 100μm. (D) mRNA expression of collagen I, III and IV as well as fibronectin was increased in Alb-cKO livers compared to CT. N=13/21 for collagen I, 10/10 for collagen III, 10/10 for collagen IV, 8/9 for fibronectin. *p<0.05. (E) Collagen is increased in the liver of Alb-cKO animals. Collagen content was evaluated using a biochemical method to quantify hydroxyproline followed by adjustment to collagen amount. N=18/21, ***p<0.001. (F) Collagen I is increased in Alb-cKO livers by western blotting. N=8/10. (G) An increase in collagen I is suggested by immunofluorescence staining. Bars represent 100μm. (H) Despite the increase in matrix, no evidence for liver-related blood laboratory abnormalities. N=31/40. Livers and blood was examined in 14-16-week-old animals, while hepatocytes were isolated by liver perfusion from 8-10 week-old animals and examined immediately. Data were analyzed using unpaired t-tests for all graphs presented in this figure. In the case of collagen IV mRNA expression Welch’s correction was applied because of the significant difference in the variances between CT and Alb-cKO. All graphs show CT to the left and Alb-cKO to the right.

We evaluated liver histology and liver function in 16-week-old animals. Sirius-red and trichrome staining confirmed increased matrix amount in Alb-cKO livers (Figure 1C). Furthermore, mRNA expression of collagen type I, III, IV as well as fibronectin was elevated (1D), as was collagen protein as determined by hydroxyproline assay, western blotting (1E-F), and staining (1G). Surprisingly, peripheral blood parameters of hepatocyte injury and liver function were not affected (1H).

Taken together, these findings suggest that loss of β1 integrin in hepatocytes increases extracellular matrix accumulation, in particular collagen I.

### Deletion of β1 integrin is associated with increased TGF-β signaling and myofibroblast activation

A possible mediator for excess matrix production and the development of fibrosis is TGF-β. In line with hepatocytes being a major source of TGF-β ^19^, TGF-β mRNA and the readily available active TGF-β protein were higher in the liver of Alb-cKO animals, while total TGF-β protein, which includes the TGF-β pool in the matrix, remained unchanged (Figure 2A). Increased TGF-β resulted in enhanced signaling as confirmed by elevated phosphorylated SMAD2, 3, and 4, while the inhibitory SMAD7 was diminished (Figure 2B-C). Similar to the livers, isolated hepatocytes from Alb-cKO animals showed increased TGF-β mRNA and protein (Figure 2D).

**Figure 2.**
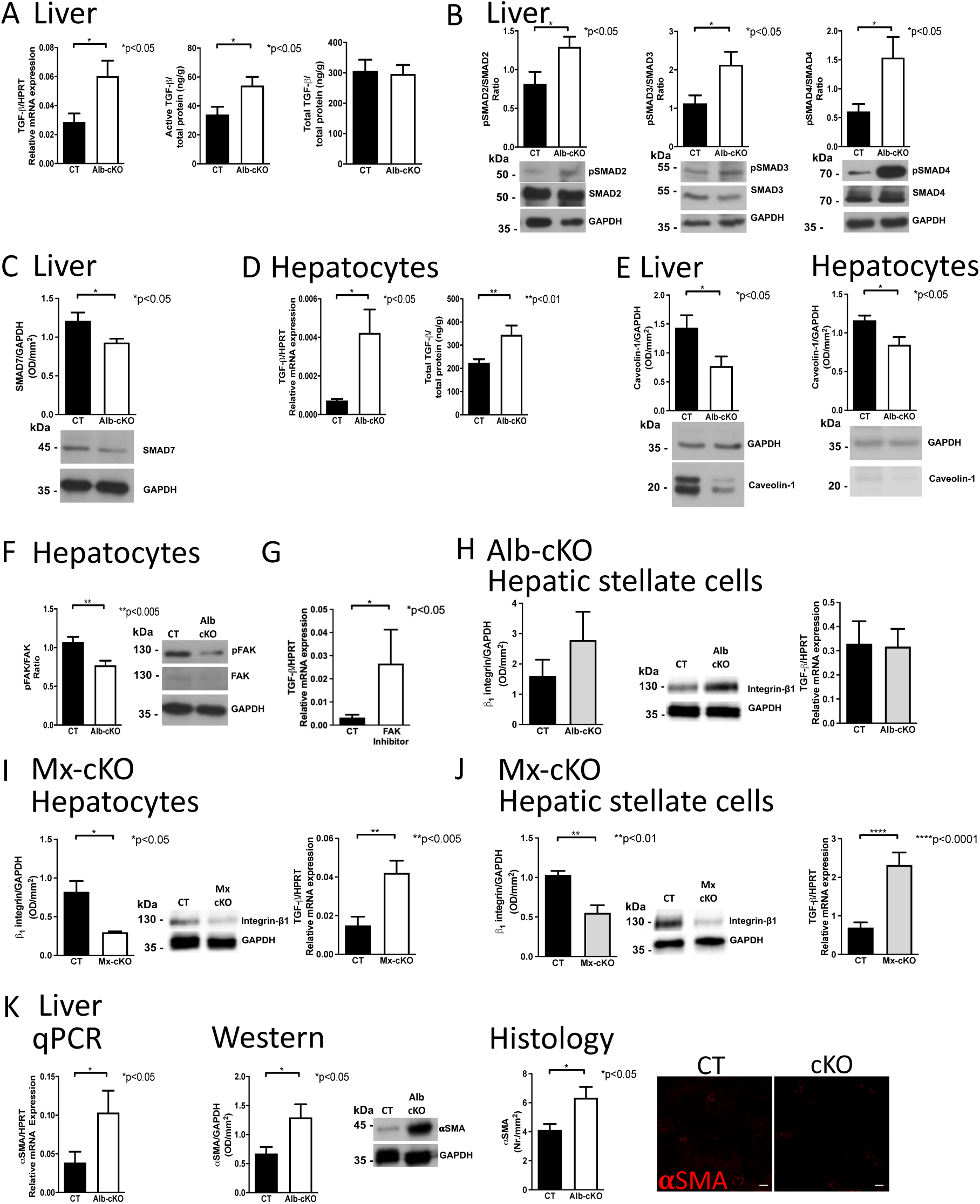
Deletion of β*1 integrin increases TGF-*β *and subsequently activation of myofibroblasts*. (A) TGF-β mRNA and active TGF-β, but not total TGF-β (which includes the matrix-bound TGF-β) were increased in livers from Alb-cKO animals. N=22/25 for mRNA, 20/19 for TGF-β protein. *p<0.05. (B-C) TGF-β signaling is enhanced in Alb-cKO livers as evidenced by western blot analysis showing increased pSMAD2, 3, 4 (in C) and decrease in the inhibitor SMAD7 (in C). N=8/9 for pSMAD2, 10/12 for pSMAD3, 4/4 for pSMAD4, and 15/20 for SMAD7. *p<0.05. (D) Evaluation of Alb-cKO hepatocytes confirms the increase in TGF-β mRNA and protein. N=6/6 for mRNA, 45/35 for active TGF-β and 16/15 for total TGF-β. *p<0.05, **p<0.01. (E) Caveolin-1 is diminished both in Alb-cKO livers and hepatocytes. N=11/14 for the livers and 7/8 for the hepatocytes. *p<0.05. (F) Isolated hepatocytes maintained for 45 min without serum show a decrease in baseline pFAK expression compared to control (CT) hepatocytes. N=16/15. **p<0.005. (G) Inhibition of FAK in cultured wildtype hepatocytes leads to a rise in TGF-β mRNA expression. Hepatocytes were cultured on vitronectin and treated for 24 hours with 5μM of the inhibitor PF573228 (Tocris). N=9/4. *p<0.05. (H) Hepatic stellate cells from Alb-cKO animals show no decrease in β1 integrin, and no change in TGF-β mRNA expression. N=7/7 for β1 integrin, N=9/16 for TGF-β. *p<0.05. (I-J) In Mx-cKO animals, β1 integrin is deleted in hepatocytes (I) and in hepatic stellate cells (J), TGF-β mRNA expression is increased in both cell types. Western for β1 integrin: N=3/3 for hepatocytes and 4/4 for hepatic stellate cells, qPCR: N=22/28 for hepatocytes and 12/11 for stellate cells. **p<0.005, ****p<0.0001. (K) mRNA and protein expression of α-SMA were increased in livers from Alb-cKO animals compared to littermate controls. More αSMA-stained cells were found in Alb-cKO liver sections. Bars represent 100μm. N=20/19 for mRNA, 14/14 for protein and 19/20 for histology sections. Livers were obtained from 16-week-old animals, hepatocytes were isolated from 8-10-week-old animals and hepatic stellate cells from 20-24-week-old mice. Data were analyzed using unpaired t-tests.

One reason for increased TGF-β is hampered caveolin-mediated internalization and subsequent diminished degradation of TGF-β ^20^. Caveolin-1 expression was lower both in the liver and in isolated hepatocytes from Alb-cKO animals (Figure 2E), consequently, TGF-β was increased ^19^.

One of the early molecules that can become phosphorylated once integrins are bound to a ligand is focal adhesion kinase (FAK). Baseline pFAK expression was suppressed in isolated Alb-cKO hepatocytes consistent with decreased integrin signaling in the absence of β1 integrin (Figure 2F). In line with the relevance of this signal, inhibiting FAK pharmacologically, and hence integrin-mediated signaling in wildtype hepatocytes increased TGF-β mRNA expression (Figure 2G). This confirms that loss of β1 integrin or FAK signaling leads to elevated TGF-β.

To determine whether depletion of β1 integrin in other cell types also affects TGF-β, hepatic stellate cells, which produce matrix in liver fibrosis, were investigated. In Alb-cKO where β1 is not depleted in hepatic stellate cells, TGF-β mRNA expression was unchanged in myofibroblasts (Figure 2H). In contrast, deletion of β1 in both hepatocytes and stellate cells using Mx to drive cre expression, which successfully depletes β1 in both cell types, increased TGF-β mRNA in both hepatocytes and hepatic stellate cells (Figure 2I-J). Since the increase in TGF-β mRNA in stellate cells was not seen in Alb-cKO mice (Figure 2H), we conclude that depletion of β1 in stellate cells, similarly to hepatocytes, leads to enhanced TGF-β mRNA expression.

In fibrosis, excess matrix is produced primarily by activated myofibroblasts that express α-smooth muscle actin (αSMA). In Alb-cKO livers, αSMA mRNA and protein expression, as well as the number of αSMA-stained cells were all increased (Figure 2K). This is consistent with enhanced myofibroblast activation in the absence of β1 integrin in hepatocytes, likely due to increased TGF-β.

In summary, loss of integrin-mediated signaling in hepatocytes results in enhanced TGF-β mRNA expression and diminished degradation, and consequently increased TGF-β protein. TGF-β then perpetuates excessive matrix production and fibrosis.

### Identification of the mediating integrin pair and *in vitro* evaluation of candidates to modulate TGF-β in hepatocytes

Deletion of β1 integrin (Figure 1) or inhibition of integrin signaling (Figure 2F) increased TGF-β and indirectly matrix production. β1 integrin can pair with a variety of α subunits to bind different matrix proteins. To determine which α subunit pairs with β1 integrin to mediate TGF-β changes, isolated hepatocytes were cultured on three matrices: collagen I, fibronectin and vitronectin. After 24 hours, TGF-β in the wells was measured. Only collagen I was associated with decreased TGF-β (Figure 3A), suggesting that engaging a collagen receptor suppresses TGF-β. We therefore evaluated the expression of all 4 collagen-binding α subunits on hepatocytes. We also analyzed α5 and αv because α5β1 and αvβ1 represent the classical fibronectin-binding integrin and a vitronectin receptor respectively. α1 and α11 expression was higher than the other two collagen-binding subunits (Figure 3B).

**Figure 3.**
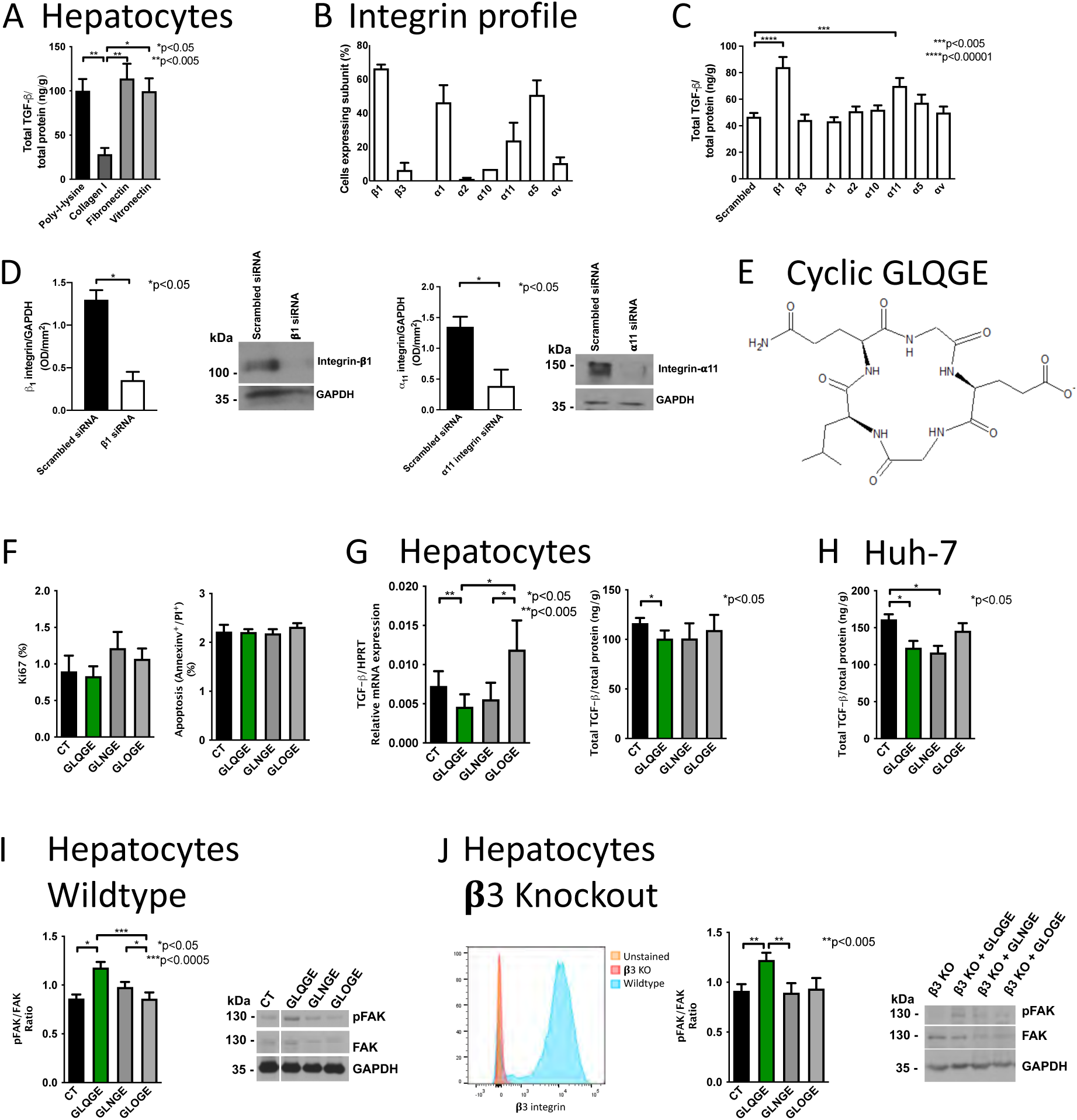
*Modulation of* β*1 integrin in vitro.* (A) Total TGF-β protein decreased when hepatocytes were cultured on collagen type I, compared to cells cultured on fibronectin or vitronectin. Freshly isolated hepatocytes were cultured in wells precoated with poly-L-lysine 0.01% or the matrix proteins at a concentration of 10μg/mL for 24 hours. Total TGF-β protein was evaluated in the medium and cells and corrected to protein content. N=11/6/11/13 replicates. *p<0.05. (B) Integrin subunits expression on the surface of freshly isolated hepatocytes was evaluated by flow cytometry. N= 3 experiments. (C-D) Knockdown of β1 or α11 integrin using siRNA in hepatocytes increased total TGF-β protein (C). Depletion was confirmed by western blotting (D). Isolated cells were cultured and treated with siRNA targeting integrin subunits that bind to collagen, fibronectin and vitronectin. TGF-β: N=36/36/21/25/33/25/25/25/25 replicates in the order of the bars, β1 depletion: N=2/2, α11 depletion N=4/2. (E) Structure of cyclic GLQGE. (F) GLQGE did not affect hepatocyte proliferation (ki67 staining, left graph) or apoptosis (annexinv-propidium-iodide staining, right graph). N=7/8/7/7 for proliferation and 7/8/8/8 for apoptosis. Cells were cultured 24 hours in the presence of 50μg/mL of the molecules, stained and evaluated by flow cytometry. (G) In hepatocytes, only GLQGE diminishes TGF-β mRNA and protein compared to control (CT). Hepatocytes cultured on vitronectin were treated with 50μg/mL of the peptides and evaluated 24 hours later. N=10/9/10/10. *p<0.05. (H) TGF-β protein in the Huh-7 hepatoma cell line was reduced after GLQGE or GLNGE treatment. Huh7 cells were cultured on vitronectin, and treated for 24 hours with 50μg/mL of the molecules. N= 5 experiments with 2-9 replicates per experiment. *p<0.05. (I) pFAK is increased after treatment of hepatocytes with GLQGE. Cells were cultured in suspension without FCS for 2 hours and treated with 50μg/mL of the peptides for 30 minutes. N=35/29/30/30. *p<0.05, ***p<0.0005. (J) Increased pFAK in hepatocytes does not require β3 integrin expression. β3 hepatocytes were obtained from global β3 knockout mice (deletion confirmed using flow cytometry in peripheral blood and shown) and compared to wildtype littermate controls. Cells isolated were treated as in I. N=4/5/5/5. **p<0.01. Comparisons were performed using unpaired t-tests.

siRNA directed against various integrin subunits in hepatocytes was used. Only α11 and β1 integrin depletion increased TGF-β protein (Figure 3C-D). Thus, only α11β1 integrin suppresses TGF-β in hepatocytes.

We next hypothesized that engaging α11β1 integrin might suppress TGF-β. The peptide GLOGE (glycine-leucine-hydroxyproline-glycine-glutamic acid) attaches to collagen-binding integrins^21^. Several modifications were made, and a total of three molecules with the sequences GL**<u>Q</u>**GE (Q: glutamine), GL**<u>N</u>**GE (N: asparagine), and GL**<u>O</u>**GE (O: hydroxyproline) were pursued. The structure of GLQGE is shown in Figure 3E. Changes in cell proliferation or apoptosis in the presence of the molecules were excluded *in vitro* (Figure 3F). As shown, TGF-β mRNA expression in cultured hepatocytes diminished with treatment with GLQGE compared to untreated control. GLNGE also diminished TGF-β mRNA compared to GLOGE. Total TGF-β protein in hepatocytes, however, was only suppressed with GLQGE (Figure 3G). In the Huh-7 hepatoma cell line, GLQGE also successfully suppressed TGF-β protein, together with GLNGE (Figure 3H). Finally, only GLQGE increased pFAK (Figure 3I). Since hepatocytes isolated from β3 global knockout mice ^22^ similarly exhibited an increase in pFAK only after GLQGE treatment, we conclude that β3 integrins do not mediate this effect (Figure 3J).

Taken together, GLQGE increases integrin signaling and diminishes TGF-β in hepatocytes.

### GLQGE inhibits fibroblast activation

To further characterize the effects of GLQGE, we evaluated its role in the NIH3T3 fibroblastic cell line. Unlike in hepatocytes, TGF-β mRNA did not change after GLQGE addition (Figure 4A). Instead, αSMA mRNA and protein and consequently, collagen I diminished in response to GLQGE but not the other two molecules (Figure 4B-C). Since αSMA increases in fibroblasts upon activation and is associated with increased matrix production ^23^, this suggests that GLQGE suppresses fibroblast activation. A dose-response analysis confirmed that GLQGE inhibits fibroblast αSMA and collagen I mRNA expression over a large range of concentrations (Figure 4D). A similar effect was seen in primary hepatic stellate cells, which transition to activated myofibroblasts and produce matrix in response to injury in the liver. Isolated stellate cells showed a higher sensitivity to GLQGE than NIH3T3 cells with suppression of αSMA at 0.1 instead of 1 μg/ml GLQGE in NIH3T3 cells (Figure 4E). In these cells, suppression of protein expression of αSMA and collagen I after exposure to GLQGE was confirmed (Figure 4F).

**Figure 4.**
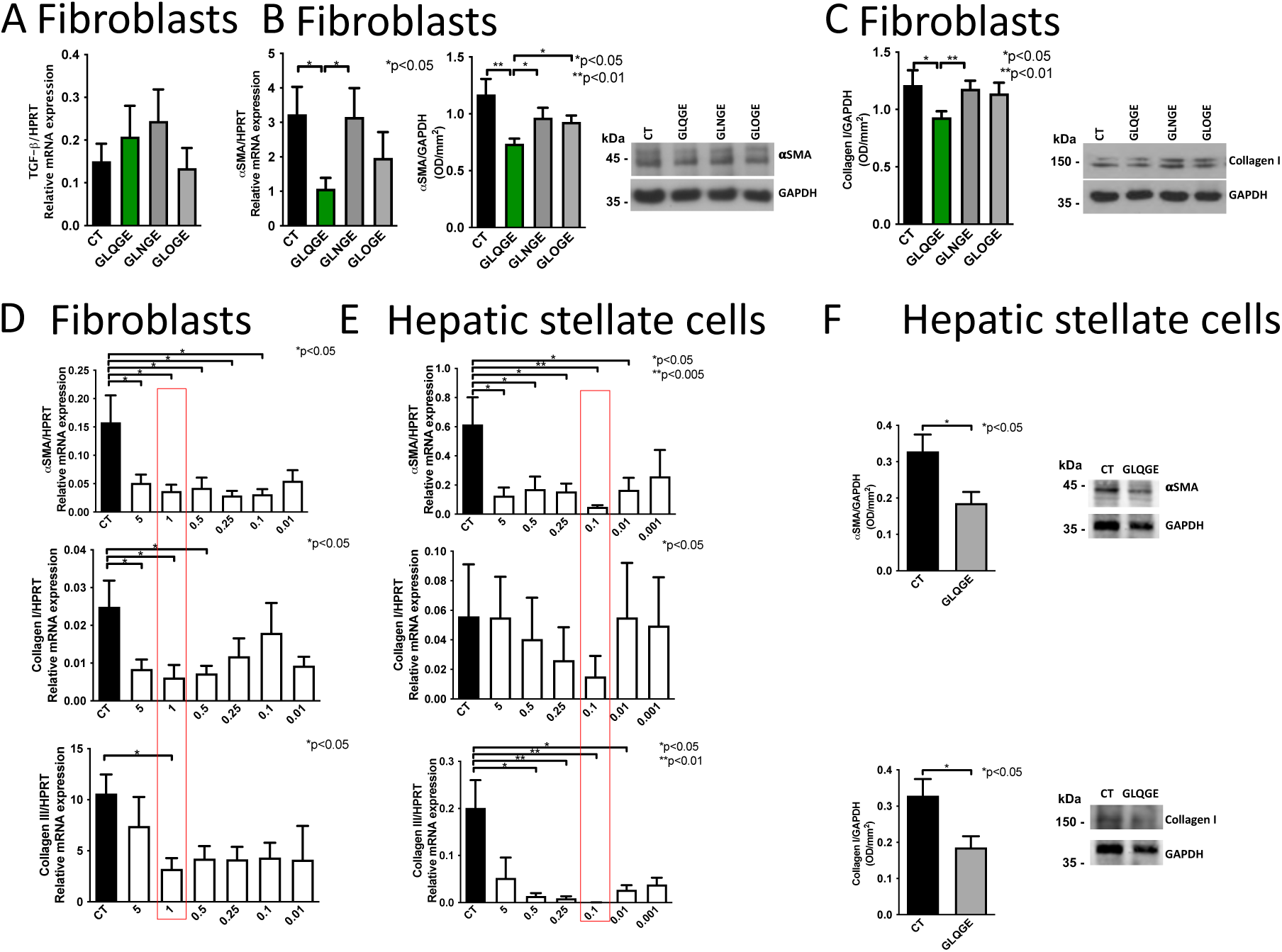
GLQGE suppresses activation of fibroblasts in vitro. (A) GLQGE does not diminish TGF-β mRNA expression in fibroblasts. NIH3T3 cells were cultured for 24 hours in the presence of the peptides at 50μg/mL. N=13/12/8/4. (B) αSMA mRNA (examined by qPCR) and protein expression (evaluated by western blotting) are decreased after GLQGE treatment. N=12/8/7/8 for mRNA and N=11/12/11/11 for protein. (C) GLQGE diminishes protein expression of collagen I in fibroblastic cells. NIH3T3 cells were cultured for 24 hours in the presence of 50μg/mL of the peptides and 10%FCS. N=60/59/56/59. *p<0.05, **p<0.01. (D) Over a broad range of concentrations (in μg/ml), GLQGE suppressed αSMA mRNA expression. Confluent NIH3T3 cells were serum-starved for 24 hours and GLQGE added in the absence of FCS for 24 hours. N=20/15/19/19/18/17/16. (E) Primary hepatic stellate cells showed suppression of αSMA mRNA and collagen III mRNA after treatment with GLQGE (in μg/ml) for 24 hours. Primary hepatic stellate cells were cultured for 24 hours after isolation with 10% FCS, then serum-starved for another 24 hours before treating with GLQGE for 24 hours. N= 8/8/10/10/10/9/8 for αSMA and 7/8/9/8/4/10/5 for collagen I. *p<0.05. (F) GLQGE diminishes αSMA and collagen type I protein expression. Primary hepatic stellate cells were cultured as in E and treated with 0.1 μg/mL GLQGE. N=10/10, *p<0.05. ANOVA followed by t-test were performed, except in D where Welch’s correction was applied because variances were significantly different. In F, unpaired t-tests were performed. CT stands for untreated cells.

Thus, GLQGE suppresses αSMA and collagen expression in a fibroblastic line and primary hepatic stellate cells *in vitro*.

### Establishing which integrins mediate GLQGE effects

GLQGE stimulated FAK phosphorylation (Figure 3I-J). Therefore, an integrin might mediate suppressed activation of fibroblasts. Since GLQGE was based on a collagen-binding sequence, we evaluated the expression of the collagen-binding integrin α subunits as well as major matrix-binding subunits on hepatic stellate cells. α1, α10, and α11 were expressed (Figure 4G). To determine which integrins bind GLQGE, we used fluorescence anisotropy. For this, the four collagen-binding integrin pairs (α1β1, α2β1, α10β1 and α11β1), as well as α5β1 the classical fibronectin receptor, αvβ1, a vitronectin receptor and αvβ3 as a negative control were tested. GLQGE only binds to α10β1 and α11β1 with moderate affinity (Figure 4H). Since the receptors were commercially obtained, we evaluated their identities by mass-spectrometry using fingerprinting and confirmed their identity (albeit with some contamination) (Supplementary Figure S1). Notably, GLQGE bound to both human and murine α11β1 with similar affinity (Figure 4H).

**Figure 4.**
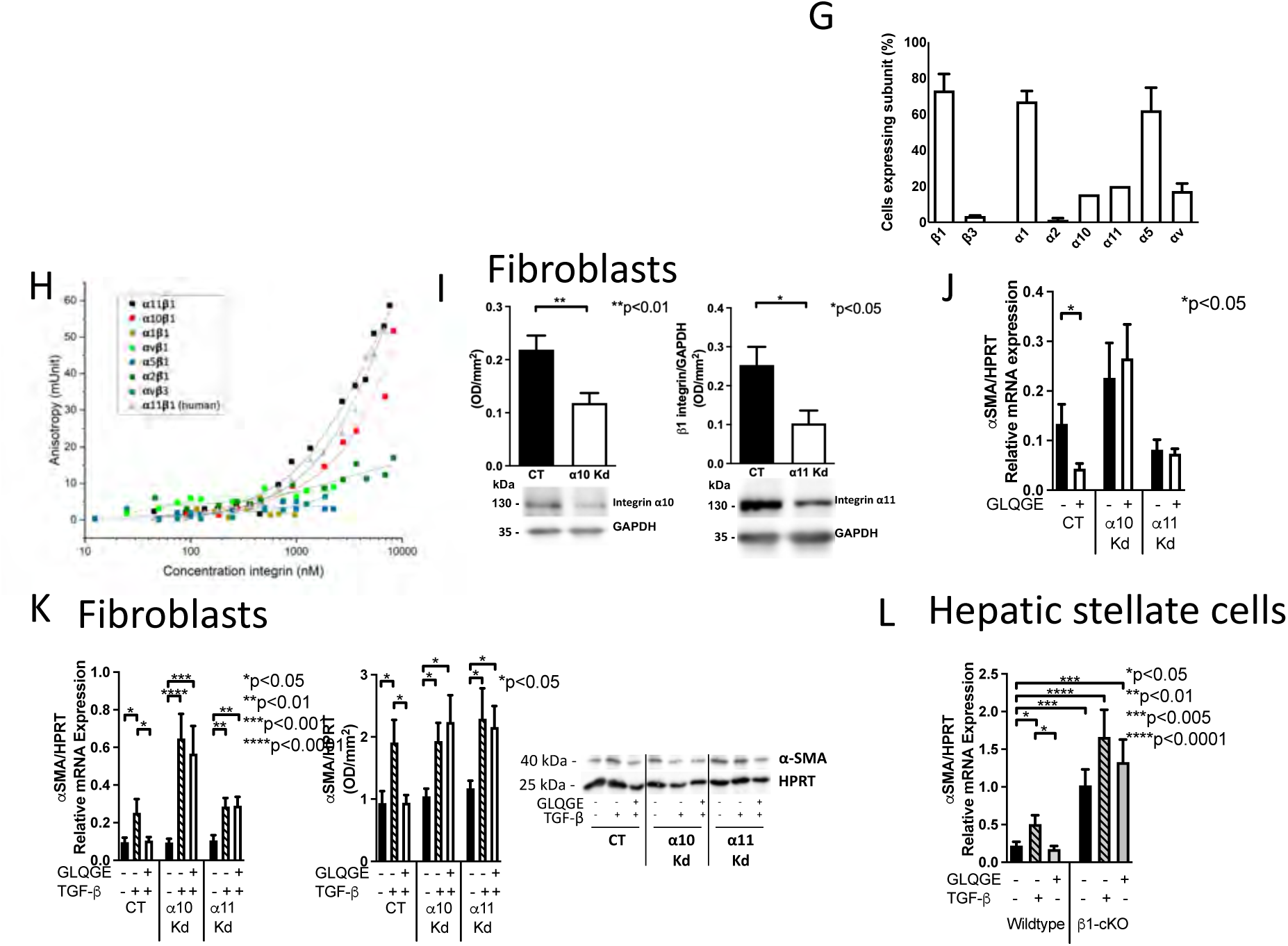
Identifying the receptor mediating GLQGE effects. (G) Hepatic stellate cells express a variety of integrin subunits including β1, β3, α1, α2, α10, α11, α5, αv evaluated by flow cytometry. N=3. (H) GLQGE binds better to α11β1 than α10β1, but not to the other integrin pairs. Fluorescence anisotropy using candidate receptors was performed in order to determine the binding affinity of fluorescently labelled GLQGE. (I) Protein expression of α10 and α11 integrin subunits is diminished in NIH3T3 after viral transduction with shRNA targeting these subunits (knockdown: Kd) compared to cells transduced with an empty vector (Control: CT). Cells were collected 24 hours after culture with FCS. N=10/11 for α10 and 4/5 for α11. *p<0.05, **p<0.01. (J) GLQGE decreases αSMA mRNA expression in control NIH3T3 fibroblasts, but not in α11 or α10 Kd cells. Cells were serum-starved for 24 hours before treatment with GLQGE 1 mg/mL for an additional 24 hours. N=27/23, 33/27, 32/31. *p<0.05. (K) TGF-β stimulation increases αSMA mRNA and protein. Combining GLQGE with TGF-β prevented this increase in CT cells, but not in α10 or α11 Kd cells. Cells were serum-starved for 24 hours, then treated with TGF-β 20ng/mL alone or in combination with GLQGE 1mg/mL for 24 hours. mRNA: N=29/22/26, 29/25/23, 28/19/20. Protein: N=14/13/12, 10/10/10, 11/11/11. *p<0.05, **p<0.01, ***p<0.001, ****p<0.0001. (L) Deletion of β1 integrin in hepatic stellate cells prevented GLQGE from restoring αSMA to baseline levels in the presence of TGF-β. β1 integrin was successfully deleted in hepatic stellate cells from Mx-cKO as shown in Figure 2J. Cells were cultured for 24 hours in FCS-containing medium, serum-starved for 24 hours, and then treated with TGF-β 20ng/mL alone or in combination with GLQGE 0.1 mg/mL. N=7/8/7, 7/8/8. *p<0.05, ***p<0.005, ****p<0.0001. Groups were compared using t-tests.

In order to confirm the relevance of the two α subunits and therefore the corresponding receptors, α10β1 and α11β1, we first knocked down (Kd) α10 and α11 in NIH3T3 cells using shRNA (Figure 4I), and confirmed that GLQGE is unable to suppress αSMA mRNA expression in α10 and α11 Kd cells compared to control (CT) (Figure 4J). αSMA increased in response to TGF-β stimulation in α10 or α11 Kd similarly to CT cells. However, while GLQGE prevented TGF-β-induced stimulation in CT, it failed to suppress αSMA in α10 or α11 Kd cells (Figure 4K). Both α10 and α11 can only bind to β1 to form an integrin dimer. We therefore took advantage of our ability to delete β1 integrin in hepatic stellate cells in Mx-cKO mice. Stellate cells isolated from these mice (depletion of β1 integrin is shown in figure 2J) responded to TGF-β, but failed to suppress αSMA mRNA in the presence of both TGF-β and GLQGE (Figure 4L).

Taken together, these data suggest that GLQGE suppresses the activation of fibroblastic cells and hence diminishes matrix production both at baseline and after TGF-β stimulation and that this requires the presence of α10β1 or α11β1 integrin on the cell surface.

### GLQGE affects MAPK and AKT pathways

To define the signaling pathways mediating GLQGE effects, we treated both hepatic stellate cells and NIH3T3 with GLQGE and performed proteomic studies. Pathway enrichment analysis revealed that GLQGE impacts ERK/MAPK ^23^ and AKT signaling^24^ pathways in both cell types (Figure 5A). A heat map of the primary cells show changes in expression of various molecules belonging to these two pathways (Figure 5B). Comparing published reports on integrin-associated proteins with the molecules that are differentially expressed in the presence of GLQGE revealed a change in ARF6^25^, MAP4K4^26^, GIT2 or LIMS1^27^ or molecules related to ras, rab- and ARFGTPases^28^ (Supplementary Table 1). We next examined the phosphorylation of two key molecules: ERK, which is part of MAPK pathway and is downregulated in response to integrin inhibition and AKT which is downstream of PI3kinase pathway and controls collagen I production. Both were downregulated in response to GLQGE treatment in control cells, but not in cells that don’t express α10β1 or α11β1 (Figure 6C-D). Thus, GLQGE inhibits AKT signaling to suppress matrix production.

**Figure 5.**
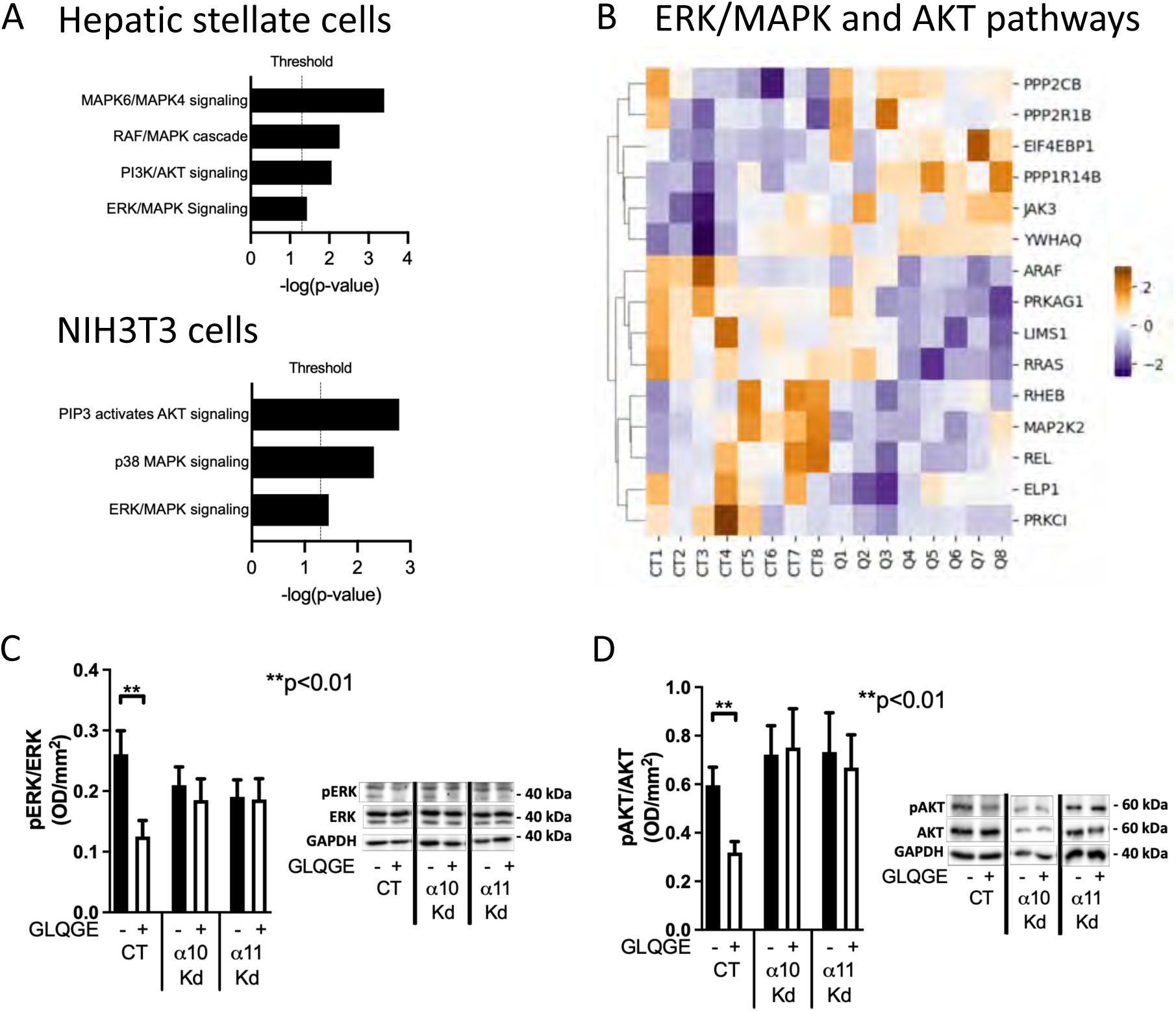
Proteomic analysis. (A-B) Pathway enrichment analysis was performed for hepatic stellate cells (treated with 0.1 μg/mL GLQGE) and NIH3T3 (treated with 1 μg/mL GLQGE) in comparison to untreated controls. A significance threshold was applied at -log(p-value) of 1.3, corresponding to a p value of 0.05. A total of 705 and 141 molecules were significantly different in t-test comparisons. Controls (CT) and GLQGE-treated (Q) cells were collected from three experiments, N= 8/8 and 6/6 replicates. (B) A heat map shows molecules associated with ERK/MAPK and AKT pathways in primary hepatic stellate cells comparing control (CT) and treated (Q) cells (with 0.1 μg/mL). N= 8/8. (C-D) Phosphorylation of ERK (C) and AKT (D) by western blotting is suppressed in the presence of GLQGE in control NIH3T3 cells, but not in α10 or α11 Kd cells and hence lacking α10β1 and α11β1 integrins. Cells were serum-starved for 24 hours, and then treated with GLQGE 1 mg/mL for 1 hour. N=10 replicates in all groups for ERK and 15/15/13/14/14/14 replicates for AKT. t-tests were performed. **p<0.01.

### GLQGE is well tolerated

Since GLQGE diminishes fibroblast activation and matrix production *in vitro,* we first compared its effects to compounds for fibrosis treatment. GLQGE was as good as BMS-986020 in its direct effects and as good as BMS or pirfenidone in counteracting TGF-β effects *in vitro* (Supplementary Figure 2A-B).

For GLQGE to be used *in vivo*, it needs to enter the circulation. Indeed, a GLQGE peak concentration in the bloodstream was measured 15 minutes after either an intraperitoneal (i.p.) injection of 0.5 mg/mouse or a subcutaneous (s.c.) injection of 1 mg/mouse. The peptide was cleared from the circulation within two hours in both cases (Figure 5E). Subsequently, we evaluated whether the cyclic peptide can reach concentrations in the liver that can be unambiguously attributed to its infiltration in the tissue, eliminating the possibility of false positive or skewed results due to peptide stemming from residual blood in the liver. At 45 minutes after either i.p. or s.c. injection of GLQGE, the amount of GLQGE per mg tissue was higher than the concentration of the molecule in the blood, suggesting that GLQGE can in fact infiltrate the tissue and its detection in the liver is not due to its presence in the circulation (Figure 5F).

**Figure 5.**
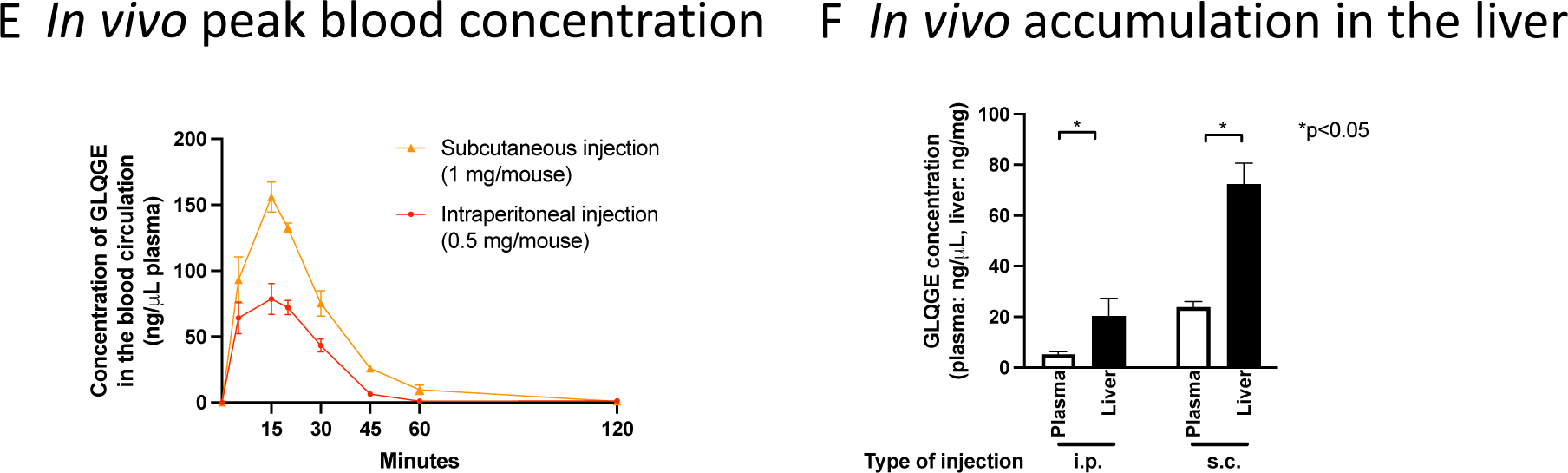
GLQGE detection in plasma and liver tissue samples. (F) GLQGE enters the circulation within 5 minutes and reaches its peak concentration after 15 minutes, both, after intraperitoneal (i.p.) injection of 0.5 mg GLQGE and after subcutaneous (s.c.) injection of 1 mg. Within 120 minutes, the peptide is cleared from the bloodstream. For i.p. injections the following number of replicates was analyzed per time point: N=4/4/4/8/6/4/3, whereas for s.c. injections N=4/4/4/8/7/4/3 replicates were measured. (F) At 45 minutes after injection, GLQGE could be detected in the liver after both injection types at a concentration higher than that in the blood stream (assuming 1 μL blood is equivalent to 1 mg tissue). N=10/10/8/8 (corresponds to the order of the samples in the bar chart). *p<0.05 by paired nonparametric t-test.

Since GLQGE can enter the circulation and infiltrate tissue, we evaluated its toxicity *in vivo*. Mice were exposed to 1 mg GLQGE injection daily. After 10 days blood was drawn and complete blood cell count as well as circulating biomarkers of liver, pancreas and kidney function were determined (Supplementary Figure 2C-F). No abnormalities could be detected.

### GLQGE diminishes liver fibrosis

GLQGE decreases TGF-β release from the hepatocytes and hence indirectly myofibroblast activation (Figure 3G). Furthermore, it directly suppresses myofibroblast activation and consequently matrix production (Figure 4F). We therefore evaluated whether these effects have any relevance *in vivo* in a model of liver fibrosis. Mice received the cytotoxin CCl_4_ which induces hepatocyte death over 6 weeks leading to myofibroblast activation and production of collagen (Figure 6A). During the last 10 days mice were injected with 0.9% NaCl or 0.5 mg of GLQGE, GLNGE or GLOGE i.p. daily in parallel to continued CCl_4_ administration.

**Figure 6.**
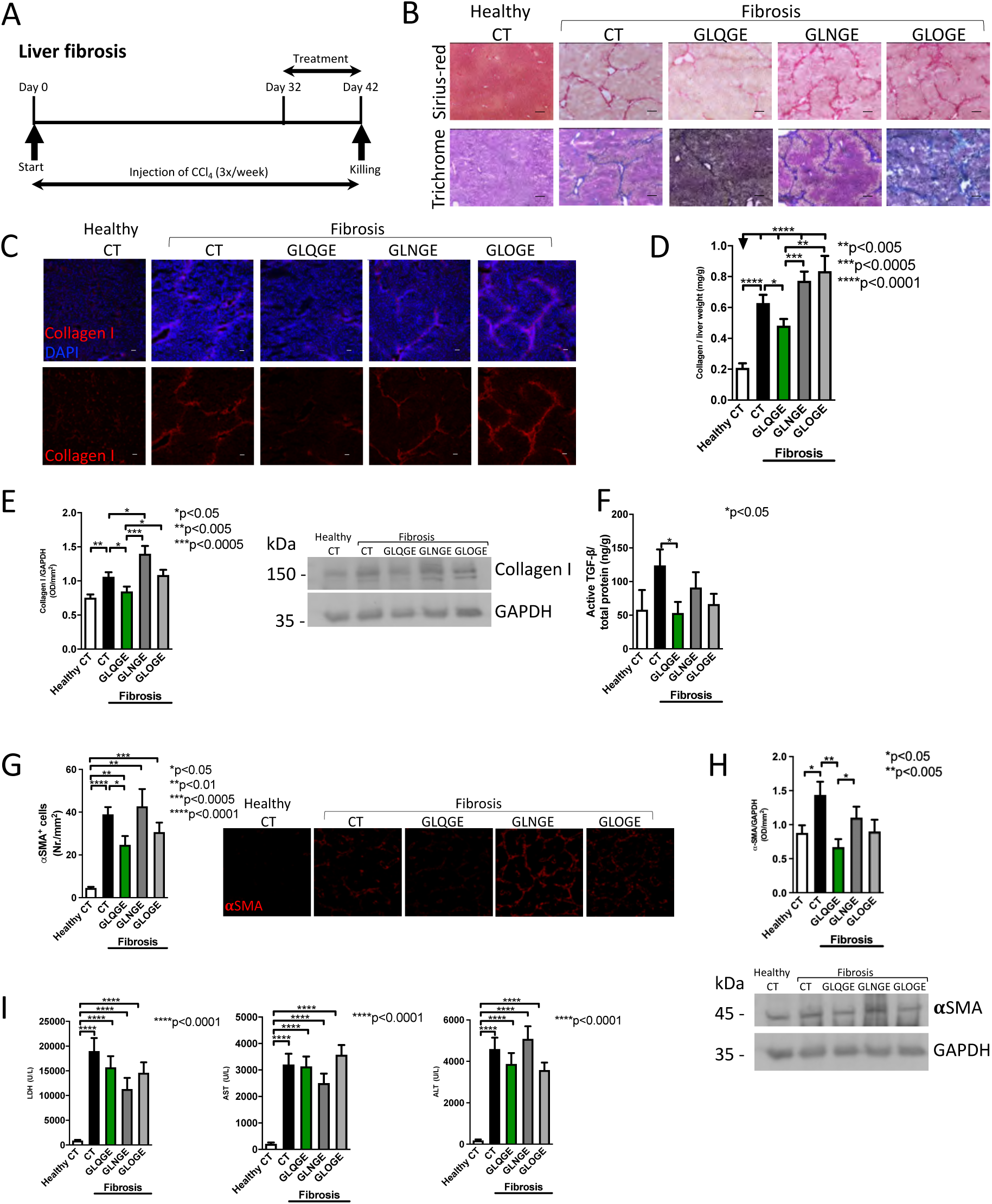
GLQGE diminishes liver fibrosis. (A) Fibrosis was induced in animals by administering CCl_4_ intraperitoneally 3x/week for 6 weeks. Over the last 10 days, animals received daily intraperitoneal injections of either 100 μL sodium chloride solution (NaCl) 0.9% (control, CT) or NaCl 0.9% containing 0.5 mg of GLQGE, GLNGE or GLOGE. Injections of CCl_4_ and the treatments were separated by at least 8 hours. (B-C) GLQGE diminishes extracellular matrix accumulation in the liver of fibrotic animals compared to fibrotic controls receiving 0.9% NaCl (Fibrosis CT), GLNGE or GLOGE as seen by Sirius-red and trichrome staining (B). GLQGE also diminished collagen (stained red) (C). Nuclei were stained for DAPI (in blue). (D-E) Collagen amount is reduced in the liver of fibrotic animals treated with GLQGE as evidenced by biochemical measurements (D) (N=15/26/19/18/16 replicates in the order of the bars), or western blot analysis (E). N=12/12/12/12/11. *p<0.05, **p<0.005, ***p<0.0005. (F) GLQGE treatment diminished active TGF-β, while total TGF-β was not affected. These results are opposite to those seen in β1 integrin Alb-cKO animals. 14/24/13/11/10 for active TGF-β and N=14/5/18/14/19 for total TGF-β. *p<0.05, ***p<0.001, ****p<0.0001. (G) The number of αSMA^+^ stained cells in the liver was diminished in GLQGE-treated animals. N=6/8/14/10/8 replicates in the order of the bars. *p<0.05, **p<0.01, ***p<0.0005, ****p<0.0001. (H) Western blot analysis confirms a reduction in αSMA protein in the GLQGE-treated fibrotic group. N=8 replicates/group. *p<0.05, **p<0.005. (I) Blood markers representing cellular injury (LDH) and hepatocyte injury (AST and ALT) showed no difference between the fibrotic groups irrespective of treatment. N=29/30/34/32/27 for LDH and AST and 29/34/32/27/27 for ALT. ****p<0.0001. Bars represent 100 μm. All numerical data in this figure were analyzed by ANOVA and if significant were followed by t-tests.

Induction of fibrosis increased matrix accumulation as evidenced by Sirius-red, trichrome and collagen I staining compared to healthy mice (Figure 7B-C). Less matrix was seen in fibrotic mice treated with GLQGE, compared to fibrotic mice injected with 0.9% NaCl, GLNGE and GLOGE. Biochemical collagen quantification and western blot analysis confirmed decreased collagen I in GLQGE-treated animals (Figure 6D-E). Active TGF-β was also diminished with GLQGE treatment (Figure 6F), raising the possibility that decreased active TGF-β might have diminished fibrosis. Both the number of αSMA^+^ cells and αSMA protein by western blotting which reflect increased myofibroblast activation diminished with GLQGE treatment (Figure 6G-H). Blood values of LDH and the two enzymes that are released in hepatocyte injury ALT and AST were increased in the blood of all fibrotic animals, but GLQGE did not lower them, suggesting that matrix reduction is not due to preventing hepatocyte toxicity (Figure 6I). Finally, the number of CD45^+^-stained immune cells was lower in GLQGE-treated livers, presumably due to diminished TGF-β (Supplementary Figure S3).

**Figure 7.**
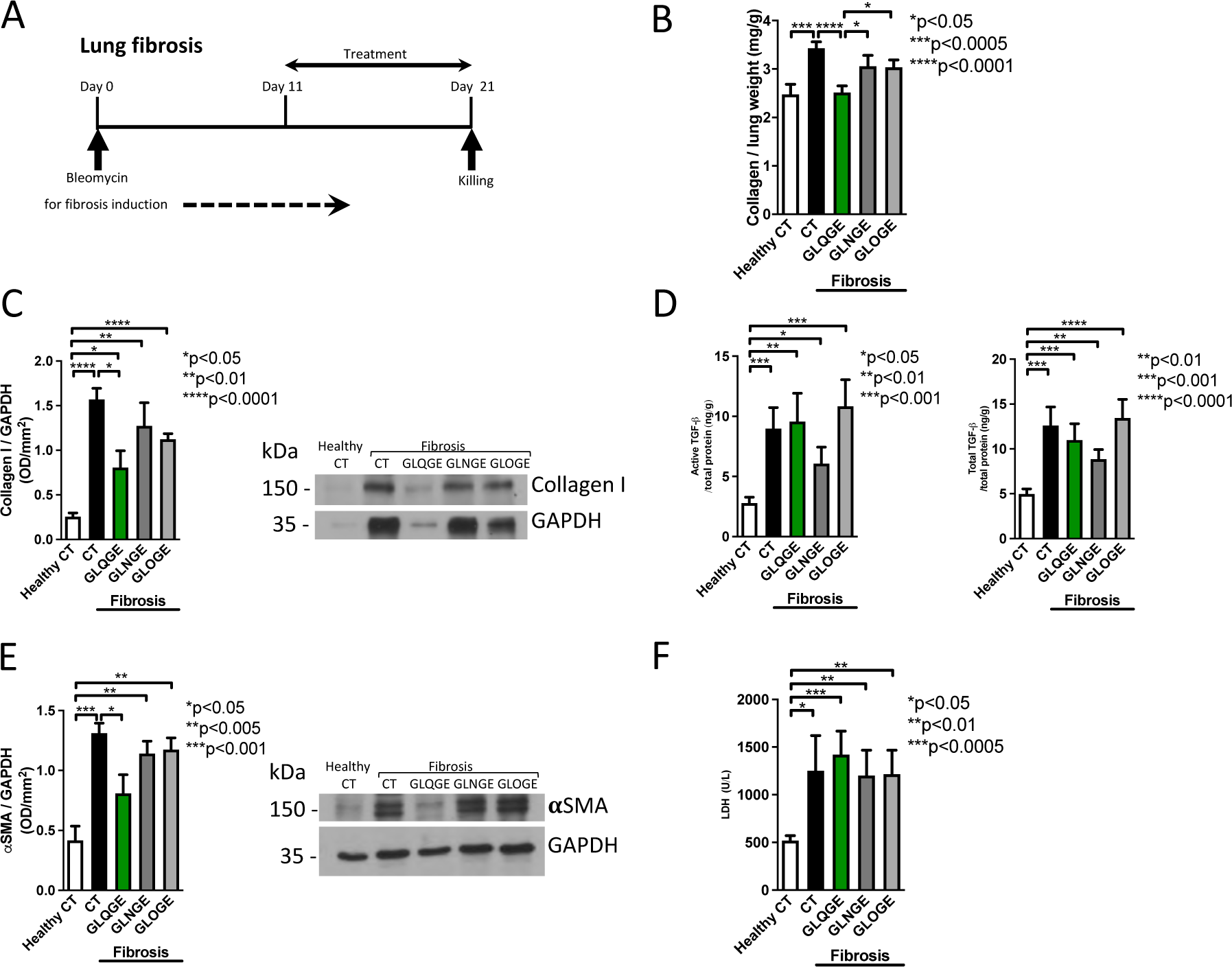
GLQGE suppresses lung fibrosis. (A) Fibrosis was induced in animals by a single intratracheal bleomycin instillation. Starting on day 11, subcutaneous injections were administered of either NaCl 0.9% (CT) or GLQGE, GLNGE or GLOGE (in NaCl 0.9%) at a dose of 1 mg/mouse/day. On day 21, the animals were euthanized. (B) GLQGE treatment diminished collagen accumulation in the lung compared to fibrotic mice receiving 0.9% NaCl, GLNGE or GLOGE. Lung lysates were evaluated biochemically. N=18/18/17/17/18 replicates in the order of the bars. *p<0.05, ***p<0.0005, ****p<0.0001. (C) Western blot analysis shows that GLQGE diminishes collagen type I in lungs of fibrotic GLQGE-treated animals compared to fibrotic NaCl-treated animals. N=4/3/3/4/4. *p<0.05, **p<0.01, ****p<0.0001. (D) GLQGE did not alter the levels of TGF-β compared to fibrotic controls. N=18/17/16/17/17. *p<0.05, **p<0.01, ***p<0.001, ****p<0.0001. (E) αSMA protein was diminished by western blot analysis of the lungs in GLQGE-treated fibrotic animals compared to NaCl-injected fibrotic CT animals. N=4/4/4/4/4 replicates. *p<0.05, **p<0.005, ***p<0.001. (F) Circulating levels of LDH, reflecting cell turnover induced by injury, were elevated in all fibrotic groups, but GLQGE treatment did not diminish LDH. N=18/18/17/17/18 replicates. *p<0.05, **p<0.01, ***p<0.0005. All numerical data were evaluated by analysis of variance (ANOVA) and when significant, t-tests were performed.

Taken together, GLQGE reduces matrix accumulation in liver fibrosis. This is associated with a lower number of activated myofibroblasts.

### GLQGE diminishes lung fibrosis

Lung fibrosis is characterized by extracellular matrix accumulation leading to compromised gas exchange. TGF-β is implicated in the pathogenesis of lung fibrosis, where activation of myofibroblasts leads to increased matrix production ^29^. We wondered whether preventing the activation of fibroblasts by GLQGE is enough to suppress lung fibrosis.

A single intratracheal bleomycin injection was applied in 8 week-old mice. Accumulation of matrix is normally detected after three weeks ^30^. Starting on day 11 GLQGE or the other two peptides were administered daily for 10 days and compared to 0.9% NaCl-treated fibrotic animals and healthy controls (Figure 7A).

Treatment with GLQGE diminished collagen detected biochemically or by western blotting in fibrotic lungs (Figure 7B-C). Sirius-red and trichrome staining similarly showed diminished extracellular matrix with GLQGE treatment (Supplementary Figure S4A-C). Interestingly, neither readily available TGF-β nor total TGF-β in the lung was diminished in GLQGE-treated fibrotic animals despite the reduction in matrix (Figure 7D). However, the number of αSMA^+^ cells (Supplementary Figure S4D) and αSMA protein were suppressed upon GLQGE exposure (Figure 7E). Similarly to the liver, the increase in circulating LDH reflecting increased cell turnover in all bleomycin-exposed animals was not affected by GLQGE suggesting that GLQGE does not protect the cells from injury and only diminishes matrix production (Figure 7F).

Taken together, these data confirm that GLQGE diminishes matrix accumulation in a second model of fibrosis, this time, without affecting TGF-β.

## Discussion

The principal finding of this work is that it is possible to diminish fibrosis by inhibiting two β1 containing integrins, namely α10β1 and α11β1 that mediate activation of fibroblasts and production of matrix. This underscores the possibility of influencing cell behavior by controlling how cells perceive their microenvironment. It also proves that the matrix is not inert or exerts only minor effects on the cells, but rather that it can modulate the severity of disease ^2^. Even though the role of the microenvironment has been the focus of much research in cancer, it has been neglected in the context of chronic diseases ^10^. This is in part a consequence of the difficulty of studying the role of matrix molecules, due to the large number of different candidates, the fact that most can bind to different receptors, sometimes simultaneously, and the extensive range of functions that can be attributed to these interactions ^5,15,16,31,32^.

β1 deletion is detrimental to various cells, where it is required for cell survival ^10,17,33^. In hepatocytes, its deletion increases TGF-β (Figure 2A, 2D). Since hepatocytes constitute a major source for TGF-β, even small changes in its production or degradation have measurable consequences ^34,35^. In the kidney, deletion of α1 and consequent loss of α1β1 signaling similarly enhanced TGF-β signaling and fibrosis ^36^. Loss of β1-integrin in hepatocytes however has additional effects. It diminishes the response to the profibrotic epidermal growth factor (EGF) and antifibrotic hepatocyte growth factor (HGF) ^17,37,38^. Interestingly, the increase in TGF-β is not a universal consequence to the modulation of β1 integrin signaling pathways. For example, even though ILK/Pinch/Parvin (IPP) mediate the crosstalk between integrins and growth factors ^39^, ILK deletion in hepatocytes did not affect TGF-β levels ^40^. Furthermore, TGF-β signaling in the kidney uses kindlin-2 without integrin involvement ^41^.

Once TGF-β is released from the cell, it is incorporated into the matrix and requires activation ^42^. Free TGF-β then enhances its own production and is degraded in part by internalization through caveolae ^14,19,20,35^. Notably, total TGF-β was neither affected in the absence of β1 integrin *in vivo*, nor after treatment with GLQGE (Figure 2B and 5F). This suggests that fibrosis in the absence of β1 integrin is related to enhancement of TGF-β activation or, more generally, a change in the turnover of active TGF-β. While fibronectin, which can bind to β1 integrin, is involved in TGF-β activation, there is contradicting evidence that β1 integrin itself (in association with α8) is required ^11,43,44^. A change in turnover is therefore more likely. This is supported by increased TGF-β mRNA in cKO livers, consistent with enhanced production, and, by lower caveolin-1 protein levels, suggestive of decreased degradation (Figure 2A and 2D) ^19,35^. For the same reason, a change in stiffness is unlikely to be responsible, because stiff matrices increase TGF-β activation and not turnover.

Our data show that cell behavior is dictated by how it perceives its environment. Thus, culturing hepatocytes on collagen decreased TGF-β (Figure 3A), while preventing this interaction by knockdown of α11 or β1 integrin increased TGF-β (Figure 3C). α11β1 is considered a major receptor for fibrillar collagens such as collagen I, and can regulate collagen turnover ^45,46^. It was detected on various fibroblastic cell types including the bone marrow stromal cells, some of which may contribute hepatic myofibroblasts to the liver ^47^. It is also expressed on hepatic stellate cells (Figure 4G). Engaging α11β1 with GLQGE suppressed not only TGF-β in the hepatocytes and consequently stellate cell activation, but also directly inhibited myofibroblast activation (Figure 3G and 4F). This dual effect diminished liver fibrosis. In the lung, GLQGE decreased αSMA (and hence fibroblast activation) without changing TGF-β (Figure 7D). The main mode of action of GLQGE in this setting is therefore the prevention of fibroblast activation and subsequently matrix production (Figure 4E-F). Importantly, the benefit by GLQGE treatment cannot be attributed to protection of the hepatocytes or other cells from injury, because enzymes such as LDH that reflect cell injury were not diminished by GLQGE treatment (Figure 7I and 7F).

In the search for antifibrotic agents that prevent matrix accumulation, one strategy has been to suppress TGF-β ^48^ with pirfenidone ^49^. Another compound, nintedanib, inhibits other growth factors ^50^, while BMS-986278 antagonizes a receptor (lysophosphatidic acid receptor 1: LPA_1_) implicated in pulmonary fibrosis ^51^. GLQGE differs from these compounds in that it directly inhibits fibroblast activation as well as matrix production even in the absence of stimulation by binding to α10β1 and/or α11β1 integrin and suppressing ERK- and AKT-mediated signaling.

In summary, GLQGE administration reduces both αSMA-expressing cells and collagen. Therefore, GLQGE offers a therapeutic possibility to prevent fibroblast activation and hence matrix formation in diseases where pathogenesis is driven or exacerbated by excess matrix production.

## Material and methods

### Mice

Animals carrying Cre-recombinase under the control of the albumin (Albumin-Cre) or the Mx promoter (Mx-Cre) were mated over two generations with β1-floxed animals (β1^fl/fl^) to produce mice in which β1 integrin was deleted in albumin-producing hepatocytes (Albumin conditional knockout for β1 integrin (Albumin-Cre_β1^fl/fl^: Alb-cKO) and in Mx-expressing cells, which includes hepatocytes and hepatic stellate cells (Mx-Cre_β1^fl/fl^: Mx-cKO). Mx was induced at 4 weeks of age using 250 μg polyinosinic-polycytidylic acid (pIpC) injected five times over 10 days^10,14^. Global β3 knockout mice were used^22^.

Liver perfusion was performed using collagenase NB4 to isolate hepatocytes and hepatic stellate cells as detailed in supplementary material and methods^14^. Liver fibrosis was induced by injecting CCl_4_ (Sigma-Aldrich) in oil (1:5 mixture) in wildtype 7-week-old mice intraperitoneally 3x/week for 6 weeks. During the last 10 days, the animals received intraperitoneal (i.p.) injections of the molecules GLQGE, GLNGE and GLOGE in 0.9%NaCl. Injections were separated by at least 8 hours from the CCl_4_ injections^13^. Lung fibrosis was induced by instilling 0.05 U/mouse of bleomycin sulfate (#B5507-15UN, Sigma-Aldrich) intratracheally once^30^. Starting day 11, the molecules were injected daily subcutaneously for 10 days until euthanasia.

Animal studies followed international including EU, national, and institutional guidelines for humane animal treatment, complied with the ARRIVE guidelines and relevant legislation, and were approved by the appropriate office for animal welfare in the state of Baden-Wuerttemberg, Germany (Regierungspraesidium Karlsruhe). The protocols used carry the following numbers: T-49/18, T-44/20, G-114/15, G-256/14, G-60/17, G-230/17, G-34/18, G-34/19, G-53/19, G-166/19, G-235/19, G-85/20, G-86/20, G-102/20, 187/20, 210/20, 276/21, 278/21.

### Cell culture

All cells were cultured in Dulbecco’s-modified-Eagle’s-medium (DMEM)/10%fetal calf serum (FCS), 50 IU/mL penicillin, and 50 μg/mL streptomycin (P/S)(PAN-Biotech). These include hepatocytes, hepatic stellate cells, NIH3T3 fibroblasts, Huh-7, MLEC (with 1% G418)(Applichem). Transfection of hepatocytes with siRNA was performed using Lipofectamine-2000 according to the manufacturer’s protocol. The used siRNAs and detailed method are given in supplementary material and methods as are the constructs and the methods for transduction of shRNA in NIH3T3.

### Flow Cytometry

Tissue was treated with collagenase NB4 (Serva, 17454). Cells were stained with the following antibodies: β1 integrin (hamster anti-mouse PE and FITC 1:100)(Biolegend, HMß1-1); β3 integrin (hamster anti-mouse PE 1:100)(Abd-Serotec, 2C9.G2(HMß3-1); α1 integrin (hamster anti-mouse PE 1:100)(BD; clone: HMα1), α2 integrin (rat anti-mouse A647 1:100)(Biolegend, DX5); α5 integrin (rat anti-mouse APC 1:100)(Biolegend; 5H10-27); αv integrin (rat anti-mouse 1:100)(Biolegend, RMV-7); integrin α10 (rat anti-mouse A647 1:100)(R&D, FAB9757R); integrin α11 (rabbit anti-mouse 1:25)(Thermoscientific, #PA5-23897); CD45 (rat anti-mouse APC-Cy7 1:400)(Biolegend; 30-F11); fixable viability stain BV450 (BD); goat anti-rabbit A647 1:100 (Abcam, ab150079).

### RNA Analysis

RNA was isolated using RNAzol (Sigma-Aldrich) and reversed transcribed with a protocol using oligo(dT) primers (25ng/μl), dNTPS (10mM), RevertAid Reverse Transcriptase (200U/μL, Fermentas), and RiboLock RNase Inhibitor (40U/μL, Fermentas). qPCR was performed using Sensifast Probe No-ROX (Bioline) and results were normalized to murine HPRT. The probes and primers used are presented in supplementary material and methods.

### Protein Analysis

Western blotting for collagen-type-I (Santa cruz, 28654), α-SMA (Sigma-Aldrich A2547), integrin α10 (Thermofisher, PA5-67829), integrin α11 (Thermo scientific PA5-23897, Bio-techne AF6498, R&D MAB4235), integrin β1 (Millipore, #MAB1997), caveolin-1 (BD 610406), pFAK/FAK (Cell signaling 3284/Millipore 06-543), pAKT/AKT (Cell signaling #9271/9272), pERK/ERK (Cell signaling 4376/9102), pSMAD2/SMAD2 (Cell signaling 3108/5339), pSMAD3/SMAD3 (Cell signaling 9520/9513), pSMAD4/SMAD4 (Abgent 3251a, Epitomics 1676-1), and pSMAD7/SMAD7 (Thermo scientific 42-0400) were performed and adjusted to GAPDH (Sigma-Aldrich) or HPRT (Invitrogen, #PA5-106984). The secondary antibodies used were goat anti-rabbit (Dianova, 111-035-045), goat anti rat (Dianova, 112-036-071) and goat anti mouse (Biorad, 170-6516).

Hydroxyproline in liver lysates was measured using a biochemical method as published^14^. Complete blood count in plasma and lactate dehydrogenase, aspartate and alanine aminotransferase, albumin and pseudocholinesterase were measured in serum obtained at the time of euthanasia using routine clinical chemistry^13^. Active and matrix TGF-β were determined in liver samples as published^14^ using a bioassay detailed in supplementary material and methods, which only detects bioavailable TGF-β, and corrected to total protein (BCA, Thermofisher).

### Peptide synthesis

Exploratory experiments were performed with linear pentamers including GLQGE, GLNGE and GLOGE, followed by cyclized peptides with proline synthesized at the Biochemistry Core Facility (RRID:SCR_025743) of the Max Planck Institute for Biochemistry. The data presented are however all with cyclic GLQGE, GLNGE or GLOGE without proline produced by BioCat GmbH and by Genosphere Biotechnologies.

### Proteomic evaluation

Cells were cultured to confluence in the presence of 10% FCS then serum-starved for 24 hours before treating with GLQGE for an additional 24 hours and taken up in lysis buffer and digested. The analysis was performed at the Max Planck Institute of Biochemistry Mass-spectrometry Facility (see supplementary material and methods)^52^. Data were finally evaluated using Ingenuity Pathway Analysis platform (RRID:SCR_008653) or STRINGDB (RRID:SCR_005223).

### Staining

Trichrome and Sirius-red staining were performed using commercial kits (Morphisto 18156 and 13425)^14^. Lung tissue was not perfused at the time of killing. Collagen I was stained using goat anti-mouse antibody (Southern Biotech #1310-01), 1:100 in PBS/5%BSA. The secondary antibody was mouse anti-goat-Cy3 (Dianova, #205-165-108). DAPI was added to stain the cell nuclei. Staining for α-smooth muscle actin (α-SMA) used an antibody coupled to Cy3 (clone 1A4, Sigma #C6198, 1:100). Pictures were made using an ECLIPSE Ti from Nikon.

### Fluorescence anisotropy

Fluorescence anisotropy measurements were carried out at 22°C on an Infinite M1000Pro (Tecan). c(GLQG[E-ed-5fam]) was used at a concentration of 11 nM. The following murine integrin pairs were purchased from R&D systems: α1β1 #8188-AB, α2β1 #7828-A2, α10β1 #7827-AB, α11β1 #7808-AB, α5β1 #7728-A5, αvβ1 # 7705-AV, αvβ3 #7889-AV, as well as human α11β1 #6357-AB. The data were analysed by nonlinear regression fitting using Bioeqs^53^. Identity of the two integrins that bound GLQGE was confirmed by peptide mass fingerprinting. More details are in supplementary material and methods.

### Pharmacokinetic studies

For evaluation of the uptake of GLQGE *in vivo*, mice were injected with the peptide either i.p. or s.c. and blood was drawn or the mice sacrificed to extract the liver and/or the lung tissue, respectively, at the specified time points. EDTA-plasma, liver or lung samples were quantified as explained in supplementary material and methods^54,55^.

### Statistical Analyses

Analyses were performed using Graphpad prism (V10). Analysis of variance were used as appropriate. If global probability values were smaller than 5%, subsequent comparisons between selected group pairs were then performed using Student’s *t*, Mann–Whitney, or Wilcoxon paired tests as appropriate. Every intervention *in vivo* was repeated at least twice in separate experiments *in vivo* and *in vitro*. Results are expressed as means ± SEM.

## Acknowledgements

We thank Prof. Reinhard Fässler for invaluable support, and Prof. Axel Roers for scientific input. We acknowledge funding by the German Research Council (DFG) (NA-400/5-1; NA-400/5-2; NA-400/7; NA-400/9) and the Max-Planck Society (M.KF.A.BIOC0001/K440). We thank Camilo Aponte for discussions, Hiba Ghura for discussions and help, Qian Jiang for help, and Monique Sterrantino and Franziska Dusi for technical support.

## CRediT authorship contribution statement

**Stefan Hamelmann:** writing – review & editing, visualization, methodology, investigation. **Fatemeh Derakhshandeh:** writing – review & editing, visualization, methodology, investigation. **Tom Wagner:** writing – review & editing, methodology, investigation. **Stephan Uebel:** writing – review & editing, visualization, validation, investigation, conceptualization. **Barbara Steigenberger:** writing – review & editing, visualization, methodology, investigation. **Claire Basquin:** writing – review & editing, visualization, methodology, investigation, conceptualization. **Sebastian Fabritz:** writing – review & editing, visualization, methodology, investigation, conceptualization. **Guido Wabnitz:** writing – review & editing, methodology, investigation. **Jutta Schroeder-Braunstein:** writing – review & editing, visualization. **Inaam A. Nakchbandi:** writing – review & editing, writing – original draft, visualization, supervision, resources, project administration, methodology, funding acquisition, conceptualization.

## Data availability statement

The data are available upon request by e-mail from the corresponding author. Please write in the subject line: “Request for raw data-2025”, and specify which data are needed in the e-mail you send. Please allow 2 weeks for response.

## Supplementary Materials

- Supplementary Figures
- Supplementary Methods
- Supplementary Table 1. Results of proteomic analysis in primary hepatic stellate cells treated with 0.1 μg/ml GLQGE for 24 hours in the absence of FCS.

