## Supplementary Data (Figures, Methods, and Table) for "MODULATION OF COLLAGEN-BINDING INTEGRINS AFFECTS FIBROBLAST ACTIVATION AND INHIBITS FIBROSIS"

### Supplementary Figures

#### Supplementary Figure S1

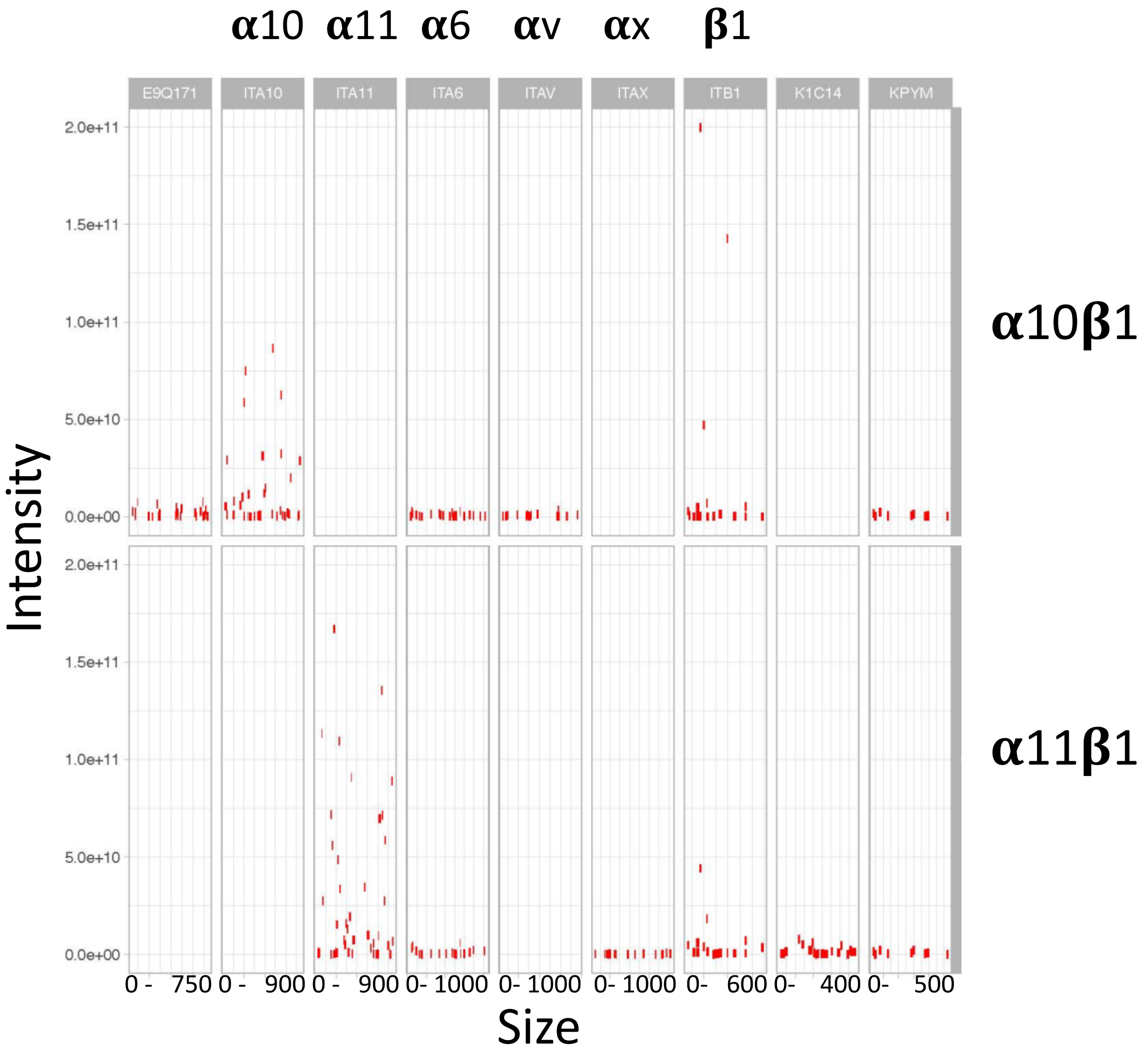


**Supplementary Figure S1**
***Finger printing***
Mass spectrometry was performed on the two murine integrins α10β1 and α11β1. The results of the two samples were plotted and evaluated by fingerprinting. α10 and β1 were identified in the upper sample (The upper sample represents α10β1) and α11 and β1 were identified in the lower sample (which represents α11β1) (Label on the right). Results for α6, αv (CD51), αx (CD11c), E9Q171 (neurofascin), K1C14 (keratin type I cytoskeletal 14), and KPYM (pyruvate kinase) are also shown.

### Supplementary Figure S2

#### Comparison with other molecules

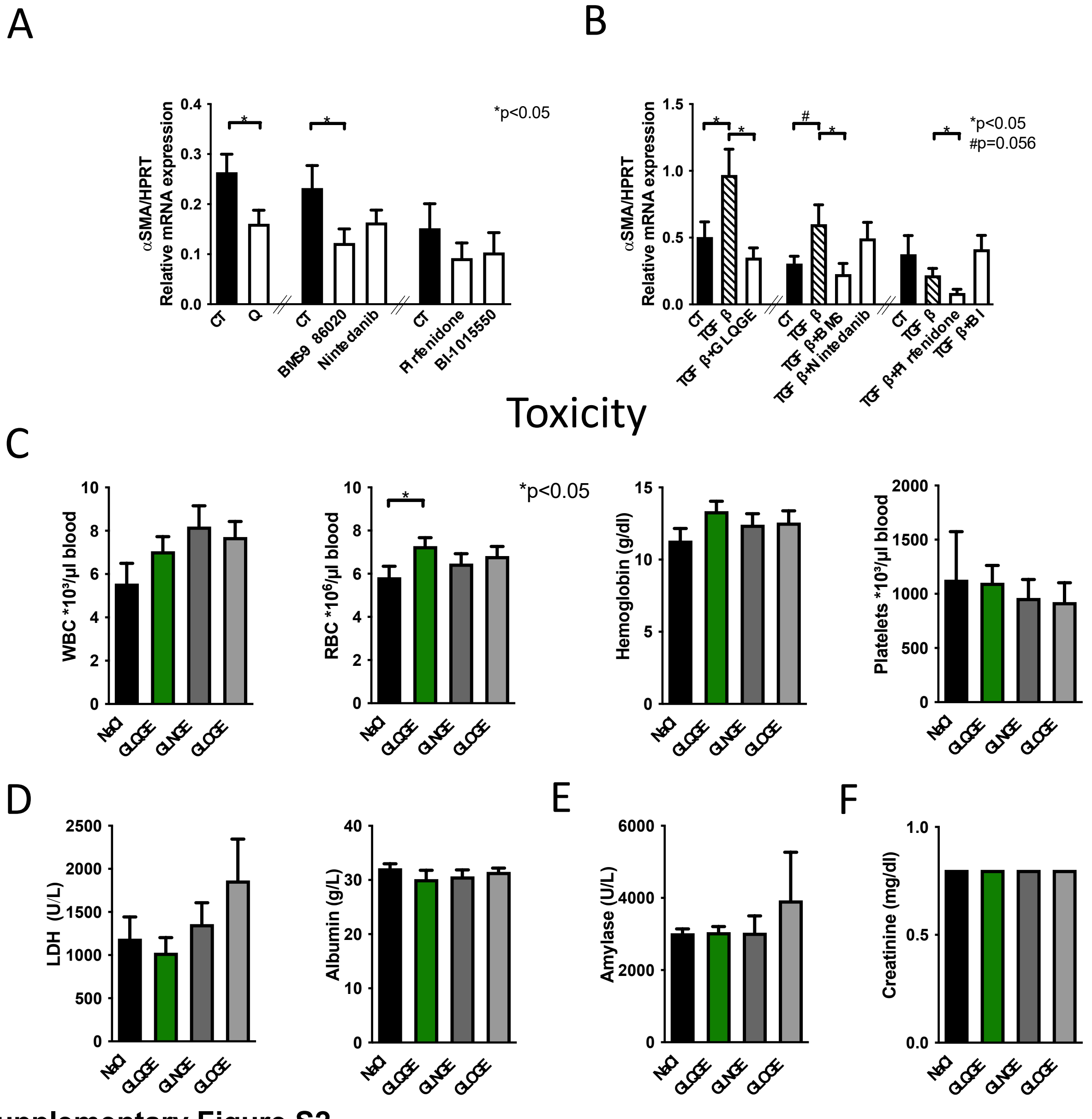


**Supplementary Figure S2.**

***(A-B) Comparison of GLQGE to anti-fibrotic molecules***

Comparing the response in NIH3T3 cells to GLQGE to 4 other commercially available substances used in fibrosis: BMS-986020, Nintedanib, BI-1015550, and Pirfenidone. Among these, only BMS-986020 diminished baseline αSMA expression. In response to TGF-β stimulation, GLQGE, BMS-986020 and pirfenidone suppressed αSMA mRNA expression. Of note, TGF-β failed to increase αSMA in the second and third groups, because of the presence of DMSO. In the second group, DMSO concentration was lower than in the third group and the increase almost reached significance. In the third group, DMSO concentration was higher. All antifibrotic molecules except GLQGE were dissolved in DMSO and the control wells received DMSO only. DMSO concentration were 12.25 mM (0.09%) for BMS-986020 and nintedanib or 122.5 mM

(0.9%) for pirfendione and BI-1015550. NIH3T3 cells were serum-starved for 24 hours, and subsequently treated with nintedanib 2 μM, BI1015550 10 μM, pirfenidone 0.5 mg/mL, or BMS-986020 10 μM in DMSO for 24 hours. Comparisons were performed using t-tests. *p<0.05, #p=0.056. N=18/19/20/21/22/18/23/22 (A) and 16/15/12/16/14/16/12/12/15/14/17 (B).

***(C-F) Evaluation of toxicity in vivo.***
Assessment of potential toxicity following 10 days of intraperitoneal administration of GLQGE, GLNGE, and GLOGE (1 mg/mouse/day) revealed no significant abnormalities in complete blood count (C) and blood values of liver (D), pancreas (E) and kidney function (F). N=10/group.

#### Supplementary Figure S3

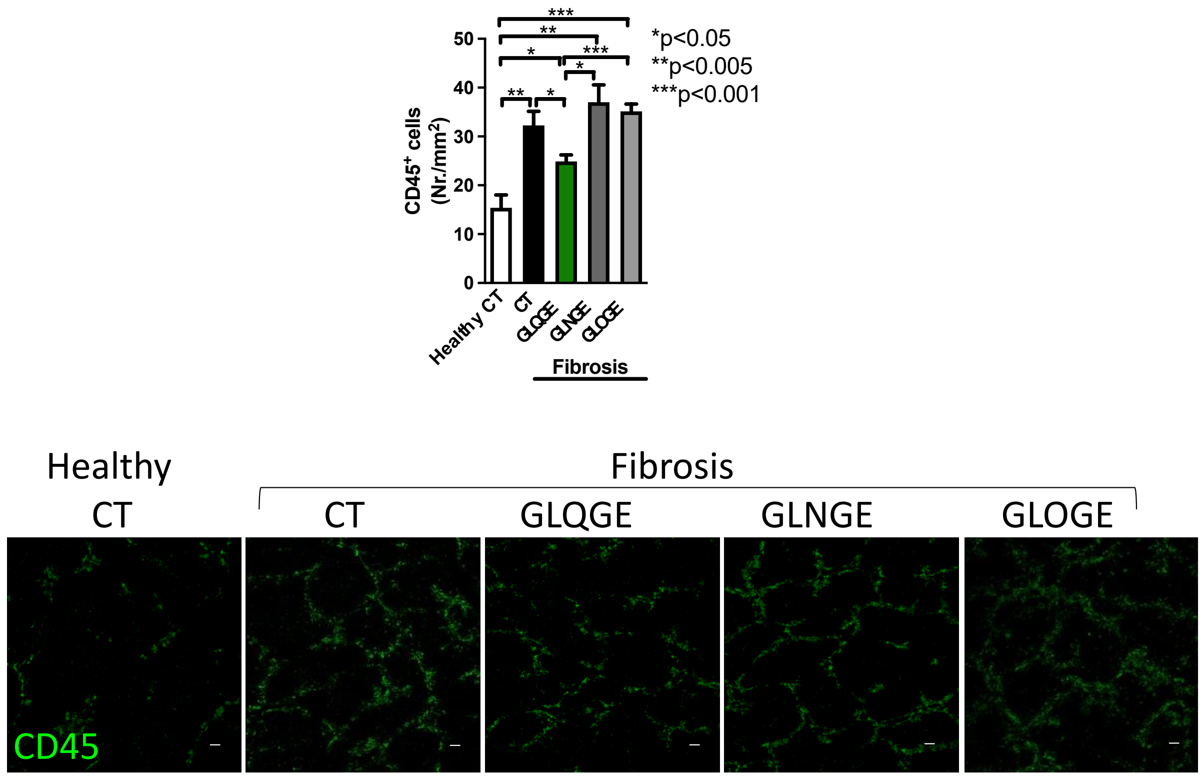


**Supplementary Figure S3.**
***GLQGE results in a decrease in CD45⁺ cells in fibrotic livers.***
Immunohistochemistry of liver sections demonstrated that the number of $CD45^+$ immune cells was increased in fibrosis, but that GLQGE was able to diminish their numbers. N=4/5/5/5/5 replicates in the order of the bars. Bars in the pictures represent 100 μm. Analysis of variance was followed by t-tests. *p<0.05, **p<0.005, ***p<0.001. The primary antibody used was a monoclonal rat CD45 antibody (Clone 30F11, BD Pharmingen, #550539), at a dilution of 1:100, and the secondary antibody used was goat anti-rat-cy2 (Dianova, #112-225-003) at a dilution of 1:500.

#### Supplementary Figure S4

##### *In vivo*
##### *Lung*

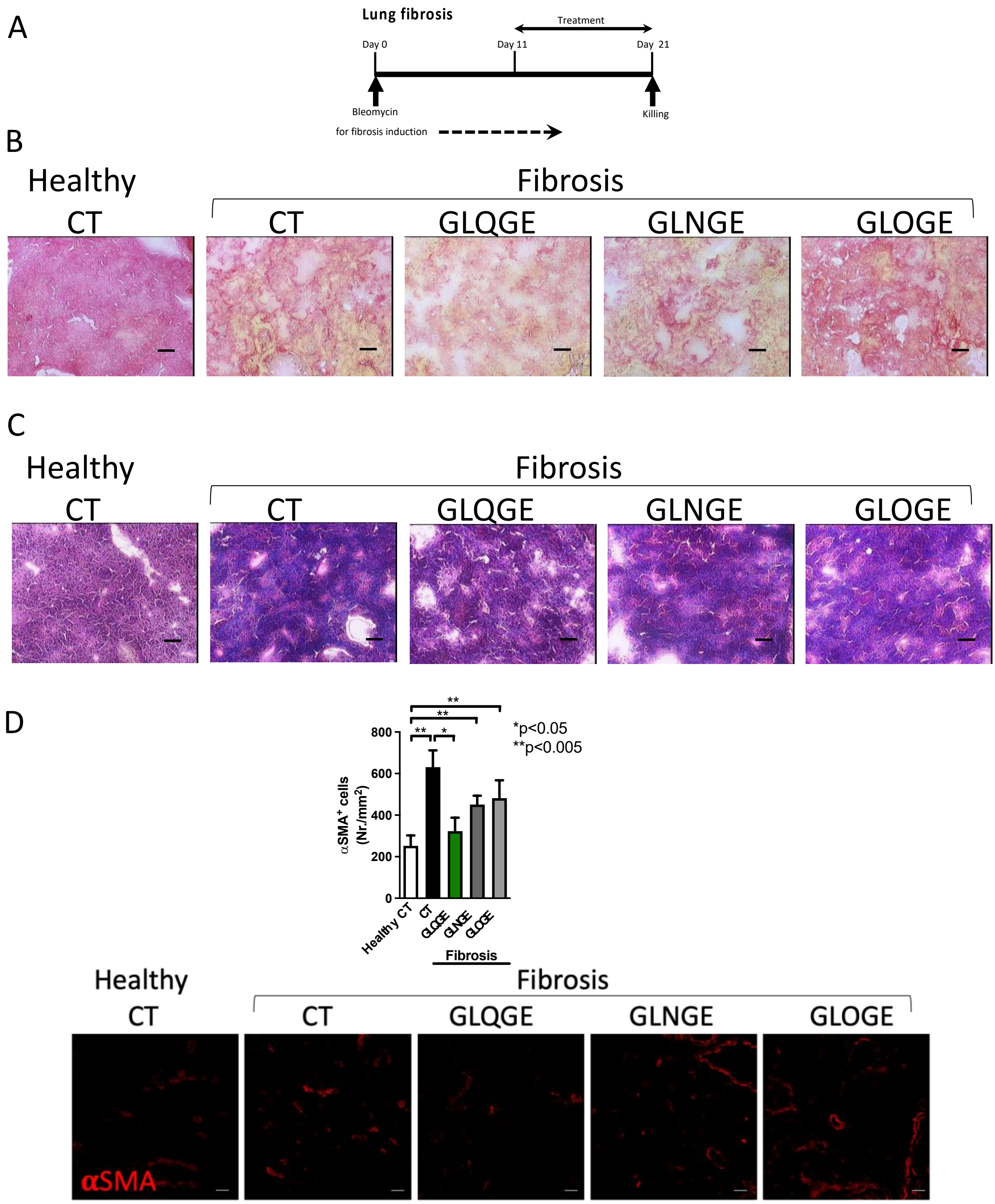


**Supplementary Figure S4.**
***Effects of GLQGE on the histology of lung fibrosis***
(A) Fibrosis was induced in animals by a single intratracheal instillation of bleomycin. Starting on day 11, subcutaneous injection with NaCl 0.9% (control, CT) or GLQGE, GLNGE or GLOGE in NaCl at a dose of 1 mg/mouse/day was administered. On day 21, the animals were euthanized. (B) Sirius-red staining shows an increase in matrix (colored red) in all fibrotic groups, but the extent was lower in the GLQGE-treated group

compared to the other fibrotic groups. (C) Trichrome staining, which colors matrix in blue suggests decreased matrix staining in the lung sections of GLQGE-treated fibrotic mice. (D) The total number of αSMA-stained cells was reduced in fibrotic mice that received GLQGE treatment. N=4/5/5/5/5 replicates. *p<0.05, **p<0.005, ***p<0.0005. Analysis of variance was followed by t-test because it was significant. In view of evidence that of the fibroblastic cells detected in the interstitium of fibrotic lungs, αSMA-stained cells represent only a subset of collagen-producing cells [31], these findings might be misleading.

### Supplementary Methods

#### siRNA experiments in hepatocytes

Lipofectamine 2000 was used according to the manufacturer's protocol (#11668027, ThermoFisher). The cells were seeded 16-24 hours before transfection to a confluence of 70-80 % in a 48-well plate in 200 µl medium per well and cultured overnight. Two reaction mixtures were prepared: one with 12.5 µl Opti-MEM® and 2 µl Lipofectamine 2000 and a second one with 12.5 µl Opti-MEM® and 30 nM siRNA. To form lipophilic complexes that facilitate the penetration of the siRNA into the cells, the two preparations were pipetted together and incubated for 5 minutes at room temperature. The transfection mixture was added dropwise to the cells for 24 hours.

**Table M1. siRNAs used in hepatocytes**

| siRNA | 5‘ siRNA | 3‘ siRNA | Source |
|---|---|---|---|
| Integrin-α1 | CCUACUUUGGUAGCGUCUU-dTdT | AAGACGCUACCAAAGUAGG-dTdT | Sigma |
| Integrin-α2 | CAGAGUACUUCAUCAAUGU-dTdT | ACAUUGAUGAAGUACUCUG-dTdT | Sigma |
| Integrin-α4 | GCAUGAAGACCAUAAUGCU-dTdT | AGCAUUAUGGUCUUCAUGC-dTdT | Sigma |
| Integrin-α5 | GAGAUGAAGAUCUACCUCA-dTdT | UGAGGUAGAUCUUCAUCUC-dTdT | Sigma |
| Integrin-α10 | GGUAUGAGGUUCACCCUUA-dTdT | UAAGGGUGAACCUCAUACC-dTdT | Sigma |
| Integrin-α11 | GUGUAUGUCUACAACCUGA-dTdT | UCAGGUUGUAGACAUACAC-dTdT | Sigma |
| Integrin-αv | CUUCUACUGGAUAACUCA-dTdT | UGAGUUUAUCCAGUAGAAG-dTdT | Sigma |
| Integrin-β1 | GUGAAGACAUGGACGCUUA-dTdT | UAAGCGUCCAUGUCUUCAC-dTdT | Sigma |
| Integrin-β3 | CCUGUAUCGCCGUACAUGU-dTdT | ACAUGUACGGCGAUACAGG-dTdT | Sigma |
| Control | Mission siRNA Universal Negative Control #1 (SIC001-10NMOL) | | Sigma |

#### qPCR primers and probes

The primers used were initially those suggested by Roche universal probe library and probes were later replaced for αSMA by 5’-atggctctgggctctgtaaggc-3’ with 5’-Fam; 3’-Tamra and for collagen I by 5’-tcctgctcctcttaggggc-3’ with 5’-Fam; 3’-Tamra (Biomers).

**Table M2. Primers und probes used in qPCR assays**

| Gene | 5‘ Primer | 3‘ Primer | Probe # |
|---|---|---|---|
| αSMA | ggagaagcccagccagtc | agcatcatcaccagcgaag | 21 and later: 5’-atggctctgggctctgtaaggc-3’ with 5’-Fam; 3’-Tamra |
| Fibronectin | tttgctcctgcacgtgttt | ctgtgtatactggttgtaggtgtgg | 66 |
| HPRT | tcctcctcagaccgctttt | cctggttcatcatcgctaatc | 95 |

| | | | |
|---|---|---|---|
| Collagen type I | catgttcagctttgtggacct | gcagctgacttcagggatg | 15 and later: 5'-tcctgctcctcttaggggc-3' with 5'-Fam; 3'-Tamra |
| Collagen type III | tggaccccaaggtcttcc | catctgatccagggtttcca | 64 |
| Collagen typa IV | agggtccccctggtctta | ccactgagcctgtcacacc | 109 |
| TGF-β | ctgggcaccatccatgac | cagttcttctctgtggagctga | 15 |

#### TGF-β MLEC assay

To analyze the TGF-β concentration in tissues, a small piece of tissue was minced in 500 µL of MLEC cell selection medium using a homogenizer and then centrifuged at 12000 g for 10 minutes. The supernatant was used to analyze the active TGF-β. The pellet was resuspended in 500 µl of fresh selection medium, incubated at 80 °C for 10 minutes, stored on ice for 10 minutes and then centrifuged at 12000 g for 10 minutes. The supernatant was used to analyze the matrix TGF-β.
For the analysis of TGF-β concentration in cultured cells, cells were seeded to 70-80 % confluence in a 48-well plate and stimulated or inhibited depending on the experiment. The medium of the cultured cells was used for the analyses of active TGF-β. The cells were mixed with 200 µl of fresh selection medium for Mlec cells, incubated at 80 °C for 10 minutes and then stored on ice for 10 minutes. The cells were transferred to a reaction tube and centrifuged at 12000 g for 10 minutes. The supernatant was used to analyze the matrix TGF-β.
MLEC (Mink lung epithelial cells) cells stably transfected with the luciferase gene under the control of a truncated PAI-1 promoter activated by TGF-β were used(*3*). The measured value in the luciferase assay correlates with the amount of expressed luciferase and therefore also with the amount of TGF-β. For this purpose, $4x10^5$ Mlec cells per well were seeded in a 96-well plate the day before the assay with the intention that they reach confluence on the next day. The medium was aspirated and the cells were washed with 100 µL DPBS. 100 µL of the appropriately diluted cell or liver lysates were added to the cells and cultured for 5 hours. To determine the TGF-β concentration, a standard curve with 2, 1, 0.5, 0.25, 0.125, 0.0625, 0.03125, 0.015612 and 0 ng/ml TGF-β (#100-21C, recombinant human TGF-β, PeproTech) in MLEC medium was applied. After incubation, 50 µL was removed from each well and 80 µL of luciferase substrate (#E6120, ONE Glo™, Promega) was added. The concentrations calculated were then adjusted to protein content measured by the BCA method (#23227, ThermoFisher).

#### Liver perfusion

After confirmation of deep anesthesia, the abdominal cavity was opened and the vena cava and portal vein were exposed. 50 mL of a solution containing EGTA (#E3889, Sigma-Alrich) (1% penicillin/streptomycin, 1% HEPES, , and 0.25% of a solution containing 3.8 g EGTA dissolved in NaOH, pH 7.4 and diluted in water to 10 mL), was pumped into the vena cava using a 25 G winged needle. At the same time, the portal vein was cut to allow the fluid pumped into the liver to flow out. To digest the connective tissue, the liver is then perfused with 50 ml of a collagenase solution 1 (0.015 % collagenase NB4 (Nordmark, #S1745403), 1 % penicillin / streptomycin, 1% HEPES,

1% $CaCl_2$ in HBSS. After perfusion of the liver, it was removed and crushed in a collagenase solution 2 using two forceps (Like solution 1 but containing 0.001 % DNase I, and 0.005 % collagenase NB4). The cell suspension was filtered using a 100 µm cell sieve and then centrifuged at 50 g without a brake for two minutes. The supernatant containing the non-parenchymal cells was removed and collected into a new tube. The hepatocytes in the pellet were taken up in 6 mL DPBS + 4 mL percoll (Cytiva, #17089102), centrifuged at 150xg for 7 min and the pellet washed again and then cultured in DMEM+10% FCS. The non-parenchymal cells were centrifuged at 800 g for five minutes without brake. The supernatant was aspirated and the pellet resuspended in 10 mL of 30 % nycodenz solution overlaid with 4 mL HBSS (Nycodenz 60% (#31000.01, Serva) was diluted 1:1 with tricine-buffer (1.8 g Tricin (#T5816-25G, Sigma-Aldrich) in 50 mL DPBS, pH 7.4) to get 30% nycodenz). The vial was centrifuged for 35 minutes at 1400 g without brake. The hepatic stellate cells, which can be recognized as white cell bands directly below the HBSS layer, were removed and collected in a new tube. The cell pellet was washed and resuspended in medium.

**Annexinv-propidium iodide and ki67 staining**

Staining for the annexin V-propidium iodide was performed by using annexin-v conjugated to Alexa 647 (#640912 Biolegend, 1:50) in buffer containing 10mM HEPES pH 7.4, 140mM NaCl and 2.5mM $CaCl_2$ for 20 minutes. The reaction was interrupted by adding an equal volume of the buffer. Propidium iodide was added 5 min before measurement. To evaluate proliferation, cells were fixed in 1% paraformaldehyde and permeabilized with 0.1% triton X (Sigma-Aldrich). The cells were then blocked in 5% BSA (Roth) for 15 minutes before staining them with an antibody directed against Ki67 (#652404, Biolegend, RRID:AB_2561525) for 30 minutes.

**Transfection/Transduction with shRNA lentiviral plasmids/viruses**

To prepare plasmids for transfection and transduction, bacterial cultures containing the shRNA plasmids were first grown on agar. Single colonies were then transferred to liquid cultures containing 100 mL of LB media (a mixture of tryptone, NaCl, yeast extract and water) supplemented with ampicillin. The plasmid DNA was then extracted using a maxi prep kit, following the manufacturer's instructions, and quantified using a photometer.
For retroviral production, HEK293T cells were seeded at $1.2 \times 10^6$ cells per 60 cm² dish (Greiner, #628160) 48 hours before transfection. DNA transfections were performed using calcium phosphate ($CaPO_4$). A mixture of plasmids: 6.5µg VSV-G (RGB007 MD2.G), 15µg psPAX2 (RGB006), and 20µg lentiviral plasmid containing the appropriate shRNA as suggested by the commercial vendors (α11: Vectorbuilder, 21-mer shRNAs nr. 1, 2, and 3 with the sequences ACGGCATTTGGCATTGAATTT, CCGCCCTGTAGTTCAAATCAA, and GCACGGCATTTGGCATTGAAT), β1: Sigma Aldrich TRCN0000066647, shRNA sequence: GCACGATGTGATGATTTAGAA. Scrambled control: Vectorbuilder shRNA#1 with the sequence: CCTAAGGTTAAGTCGCCCTCG as well as calcium chloride ($CaCl_2$) was prepared in HBS buffer. After 5 minutes of incubation at RT, plasmid mixture was added to the HEK293T cells dropwise. 24 hours post-transfection, media was replaced with low glucose DMEM (1.0 g/L, ThermoFisher 21885-025). The supernatant containing viral particles was filtered (Millex PVDF 0.45 µm) and collected 24 hour later. For viral transduction, NIH3T3 cells were seeded in 6-well plates 24 hours prior to the start of

transduction to reach 50% confluency. A mixture of collected viral supernatant and 1 µg/mL polybrene was added to the cells. After 24 hours, the medium was replaced with fresh growth medium, and selection was carried out with puromycin (4 µg/mL) or blasticidin (25 µg/mL). Successful transduction was examined via western blot (ITGA11, R&D, MAB4235), and flowcytometry (CD29, Biolegend, 102208).

**Transduction with packaged shRNA lentiviral particles**
Lentiviral transduction for knockdown of α10 was performed using packaged viral particles obtained from VectorBuilder following the manufacturer's protocol (Vectorbuilder, shRNA#1, and 2, with the sequences CCCTGAGATAGATCCAAACAA, GCGTTGGGATAGAAGATTCTA). NIH3T3 cells were seeded in complete culture medium 24 hours before transduction to reach 50% confluency. Shortly before introducing the viral vectors, the medium was replaced with fresh medium. Lentiviral particles were then added at the desired multiplicity of infection (MOI), and the plates were gently rocked to ensure even distribution. After 24 hours of incubation at 37°C in a humidified 5% $CO_2$ incubator, the virus-containing medium was replaced with fresh complete medium, and cells were allowed to recover for an additional 24 hours. 48 hours after transduction, the medium was replaced with fresh medium + 10%FCS and 25 µg/mL blasticidin. Successful transduction was confirmed via western blot analysis (ITGA10, Thermofischer, PA5-67829).

**Table M3. shRNAs used to knockdown integrin subunits in NIH3T3 cells**

| shRNA | Sequence | source |
|---|---|---|
| Integrin α11 | ACGGCATTTGGCATTGAATTT; CCGCCCTGTAGTTCAAATCAA; GCACGGCATTTGGCATTGAAT | Vectorbuilder |
| Integrin α10 | CCCTGAGATAGATCCAAACAA; GCGTTGGGATAGAAGATTCTA | Vectorbuilder |
| Integrin β1 | GCACGATGTGATGATTTAGAA | Sigma Aldrich |
| Control scrambled | CCTAAGGTTAAGTCGCCCTCG | Vectorbuilder |

**Fluorescence anisotropy**

Fluorescence anisotropy measurements were carried out in 384-well plate at 22°C on an Infinite M1000Pro (Tecan) using 20 µL reactions volume. The peptide c(GLQG[E-ed-5fam]), where [E-ed-5fam] stand for glutamic acid-γ- ethylenediamine-5-carboxyfluorescein, was dissolved in D-PBS and used at a final concentration of 11 nM. The following integrin pairs were purchased from R&D systems: α1β1 #8188-AB, α2β1 #7828-A2, α10β1 #7827-AB, α11β1 #7808-AB, α5β1 #7728-A5, αvβ1 # 7705-AV, αvβ3 #7889-AV, as well as human α11β1 #6357-AB. 250 µg of the various integrins were dissolved with 125 µL of D-PBS as a stock solution and diluted at various concentrations in the wells for the measurements. The excitation and emission wavelengths were 485 nm and 535 nm, respectively. Each titration point was measured three times using ten reads with an integration time of 40 µs. The data were analysed by nonlinear regression fitting using Bioeqs(*53*). Identity of the two integrins that bound GLQGE was confirmed by peptide mass fingerprinting.

**Proteomic analysis**

**For fingerprinting**

Sample Preparation

Cell were lysed in 1%SDS, 100 mM TRIS, 10 mM TCEP, 40 mM chloracetamide. To the protein solution, 50 µL of SDC buffer (1% SDC, 40 mM CAA, 10 mM TCEP in 100 mM Tris, pH 8.0) was added. The samples were incubated for 20 minutes at 37°C before being diluted 1:1 with water. Proteins were digested overnight at 37°C by the addition of 0.5 µg of LysC and 1 µg of trypsin. After acidification with TFA (final concentration of 1%), peptides were loaded onto Evotips.

LC-MS/MS Data Acquisition

Peptides were eluted from the Evotips onto a 15 cm PepSep C18 column (1.5 µm, Bruker Daltonics) using the Evosep One HPLC system (Evosep). The column was maintained at 50°C, and peptide separation was achieved using the 30 SPD method. Eluted peptides were directly ionized and introduced into a timsTOF Pro mass spectrometer (Bruker) via electrospray ionization. Data acquisition on the timsTOF Pro was performed using timsControl. The mass spectrometer was operated in data-dependent PASEF mode, with one survey TIMS-MS and ten PASEF MS/MS scans per acquisition cycle. Analysis was performed within a mass scan range of 100-1700 m/z and an ion mobility range from 1/K0 = 1.6 Vs $cm^{-2}$ to 0.6 Vs $cm^{-2}$, using equal ion accumulation and ramp times in the dual TIMS analyzer (100 ms each) at a spectra rate of 9.43 Hz. Suitable precursor ions for MS/MS analysis were isolated in a window of 2 Th for m/z < 700 and 3 Th for m/z > 700 by rapidly switching the quadrupole position in sync with the elution of precursors from the TIMS device. The collision energy was lowered as a function of ion mobility, starting from 59 eV for 1/K0 = 1.6 Vs $cm^{-2}$ to 20 eV for 0.6 Vs $cm^{-2}$. Singly charged precursor ions were excluded with a polygon filter mask, and further m/z and ion mobility information was used for 'dynamic exclusion' to avoid re-sequencing of precursors that reached a 'target value' of 20,000 a.u. The ion mobility dimension was calibrated linearly using three ions from the Agilent ESI LC/MS tuning mix (m/z, 1/K0: 622.0289, 0.9848 Vs $cm^{-2}$; 922.0097, 1.1895 Vs $cm^{-2}$; 1221.9906, 1.3820 Vs $cm^{-2}$).

Data Analysis

Raw data were processed using the MaxQuant computational platform (version 2.2.0.0) (*52*) with standard settings applied for Bruker ion mobility data. Briefly, the peak list was searched against the UniProt sequences of mouse (SwissProt and TrEMBL). Cysteine carbamidomethylation, methionine oxidation, and N-terminal acetylation were set as variable modifications. Trypsin/P was set as the protease, and the digestion mode was set to "specific."

**For pathway analysis**

Sample Preparation

Lysed proteins samples were then sonicated using a Bioruptor Plus sonication system (Diogenode) for 10 cycles of 30 seconds each at high intensity. After sonication, the samples were diluted 1:1 with water and digested at 37°C for 1.5 hours with 1 µg of LysC, followed by overnight digestion at 37°C with 2 µg of trypsin (Promega). The resulting peptide mixture was acidified with trifluoroacetic acid (TFA, Merck) to a final concentration of 1%, followed by desalting using Evotips.

LC-MS/MS Analysis

Peptides were eluted from Evotips onto a 15 cm PepSep C18 column (15 cm x 15 cm, 1.5 µm, Bruker Daltonics) using the Evosep One HPLC system. The column was heated to 50°C, and peptides were separated using the 30 SPD method. Data acquisition was performed on a timsTOF Pro mass spectrometer using timsControl software. The instrument operated in data-independent (DIA) PASEF mode with a mass scan range of 100–1700 m/z and an ion mobility range of 1/K0 = 0.70 Vs $cm^{-2}$ to 1.30 Vs $cm^{-2}$. Equal ion accumulation and ramp time of 100 ms each were set in the dual TIMS analyzer, with a spectral rate of 9.52 Hz. DIA-PASEF scans were acquired in the mass range of 350.2–1199.9 Da, with an ion mobility range of 1/K0 = 0.70 Vs $cm^{-2}$ to 1.30 Vs $cm^{-2}$. The collision energy was ramped linearly from 45 eV at 1/K0 = 1.30 Vs $cm^{-2}$ to 27 eV at 1/K0 = 0.85 Vs $cm^{-2}$. A total of 42 DIA-PASEF windows were distributed to one TIMS scan each, with switching precursor isolation windows, resulting in an estimated cycle time of 2.21 seconds.

Data Analysis

Raw data were processed using Spectronaut 18.0 in directDIA+ (library-free) mode. The peak list was searched against a predicted human database from Uniprot (SwissProt and TrEMBL, downloaded in 2023) and custom FASTA sequences of bait proteins. Cysteine carbamidomethylation was set as a static modification, while methionine oxidation and N-terminal acetylation were set as variable modifications. Protein quantification across samples was performed using label-free quantification (MaxLFQ) at the MS2 level. The resulting data shown in the manuscript were evaluated using the Ingenuity Pathway Analysis platform (RRID:SCR_008653) or the STRING DB platform (RRID:SCR_005223).

#### Pharmacokinetic studies

For evaluation of the uptake of GLQGE *in vivo*, mice were injected with the peptide either i.p. or s.c. and blood was drawn or the mice sacrificed to extract the liver and/or the lung tissue, respectively, at the specified time points. EDTA-Plasma was mixed 1:1 (v/v) with an internal standard (IS) peptide GLNGE solution at 1 μg/mL , followed by an addition of 100% acetonitrile resulting in a dilution of 1:10 (v/v). After vortexing for 30 seconds and centrifugation at 10000xg for 10 minutes, the supernatant was diluted 1:10 (v/v) in 5% aqueous acetonitrile before measurement (final IS concentration 5 ng/mL). To evaluate the peptide content in the organs, approximately 50 mg liver or 20 mg lung tissue were weighed and the exact weights were recorded. Subsequently, 50 or 20 μL of the internal standard solution with a concentration of 2 μg/mL were added to the tissue, which was homogenized with a pestle for 3-5 minutes afterwards. Finally, the samples were diluted 1:10 (v/v) with 100% acetonitrile, followed by vortexing and centrifugation similarly to the procedure performed for the plasma samples. The supernatant was then diluted 1:10 (v/v) in 5% aqueous acetonitrile before measurement (final IS concentration 5 ng/mL).

GLQGE peptide quantification was performed using a Sciex QTrap 6500+ mass spectrometer (MS) hyphenated to a Shimadzu Nexera UPLC system equipped with a Supelco Titan C18 column (2.1 x 100 mm, 1.9 µm), whereby the column oven was heated to 45 °C. The reverse phase chromatography was started with a 5 µL sample injection. The UPLC was operated in gradient mode using water as solvent A and acetonitrile as solvent B, both supplemented with 0.1 % (v/v) formic acid. All solvents and additives were of LCMS grade. The following gradient was used to separate the peptides from biological matrices: 0-1.5 min 5% B, 1.5-6 min B increased to 50%, 6-6.1 min B increased to 98%, 6.1-7.5 min 98% B, 7.6-7.7 min decrease to 5% B and 7.7-9 min 5% B. A total flow rate of 550 µL/ min was applied.

Subsequently, the eluents were introduced into the mass spectrometer using the following source parameters: ESI+, CUR (curtain gas) 40, CAD (collision gas) Medium, IS (ion spray voltage) 5500, TEM (heater temperature) 450, GS1 (nebulizer gas) 80,

GS2 (heater gas) 60. The MS was operated in the multiple reaction monitoring (MRM) mode. The required MS/MS fragmentation parameters are given in Table M4.

**Table M4:** MSMS (MRM) acquisition parameters for the cyclic peptides GLQGE and GLNGE: precursor mass (Q1), fragment mass (Q3), dwell time, ID, declustering potential (DP), entrance potential (EP), collision energy (CE) and cell exit potential (CXP).

| **Q1 [Da]** | **Q3 [Da]** | **Dwell time [ms]** | **ID** | **DP [V]** | **EP [V]** | **CE [V]** | **CXP [V]** |
|---|---|---|---|---|---|---|---|
| 485.2 | 355.2 | 100 | GLQGE.1 | 70 | 10 | 25 | 18 |
| 485.2 | 187.0 | 75 | GLQGE.2 | 70 | 10 | 38 | 22 |
| 471.2 | 341.2 | 75 | IS.GLNGE.1 | 40 | 10 | 27 | 14 |
| 471.2 | 187.0 | 50 | IS.GLNGE.2 | 40 | 10 | 41 | 20 |

Data analysis and MRM peak integration were performed using the Sciex Analyst 1.7 Hotfix 3 and MultiQuant 3.0.3 Hotfix 4 software.

The measured sample values were compared to those of a standard curve that ranged from 0.25-250 ng/mL of GLQGE (*54*). The resulting peptide sample concentrations were corrected based on the applied sample dilution and for organ samples additionally based on the initial water content of the respective tissues. Hence, for plasma samples a total dilution factor of 200 was taken into account and for tissue samples the following corrections were applied: liver ((weightx0.75)+1000)/weight*0.75)*10) and lung ((weightx0.815)+1000)/weight*0.75)*10) to adjust for water content. Finally, plasma values are presented as ng (peptide) /µL (plasma), while liver values were normalized to the weight as follows (calculated dilution *(weight*0.75/1000))/weight). For lung samples, 0.75 was replaced by 0.815 (based on Table 5 in (*55*)).

**Supplementary Table 1.**
**Results of proteomic analysis in primary hepatic stellate cells treated with 0.1 mg/ml GLQGE for 24 hours in the absence of FCS.**

| Genes | Q/CT ratio | Q vs CT p-value |
|---|---|---|
| Rps23 | 0,71 | 0,0000 |
| Rpl18a | 0,80 | 0,0000 |
| Dhx15 | 0,88 | 0,0000 |
| Rpl14 | 0,76 | 0,0000 |
| Nucb1 | 1,15 | 0,0000 |
| Gm17949 | 0,29 | 0,0000 |
| Arf6 | 0,82 | 0,0000 |
| Rpl28 | 0,82 | 0,0000 |
| Sod1 | 3,95 | 0,0000 |
| Dr1 | 1,20 | 0,0001 |
| Arf1 | 0,86 | 0,0001 |
| Ap2s1 | 0,75 | 0,0001 |
| Spag7 | 1,26 | 0,0001 |
| Pdia6 | 1,14 | 0,0001 |
| Rps9 | 0,86 | 0,0001 |
| Rps20 | 0,87 | 0,0001 |
| Snap23 | 1,29 | 0,0002 |
| Fam107b | 1,30 | 0,0002 |
| Rps26 | 0,73 | 0,0002 |
| Cnpy2 | 1,18 | 0,0002 |
| Cybc1 | 0,77 | 0,0002 |
| Rps3a | 0,90 | 0,0002 |
| Rpl37a | 0,71 | 0,0002 |
| Rab32 | 0,87 | 0,0003 |
| S100a13 | 1,45 | 0,0003 |
| Rpl11 | 0,89 | 0,0003 |
| Myl12a | 1,22 | 0,0003 |
| Rpl13a | 0,77 | 0,0003 |
| Cept1 | 0,79 | 0,0003 |
| Txnl1 | 1,15 | 0,0003 |
| Gpx4 | 0,86 | 0,0003 |
| Rps27a | 1,52 | 0,0003 |
| Cuta | 10,48 | 0,0003 |
| Pigs | 0,91 | 0,0003 |
| Tmem35b | 0,69 | 0,0004 |
| Isy1 | 1,21 | 0,0004 |
| Rpl35a | 0,78 | 0,0004 |
| Prpf38a | 0,83 | 0,0004 |
| Tubb4b | 1,11 | 0,0004 |
| Ucp2 | 0,47 | 0,0004 |
| Pacsin2 | 1,12 | 0,0004 |
| Npepl1 | 1,52 | 0,0004 |
| Plscr3 | 1,54 | 0,0005 |
| Rps2 | 0,87 | 0,0005 |
| Rab6a | 0,82 | 0,0005 |
| Fhip2a | 0,89 | 0,0005 |
| Ubac1 | 1,23 | 0,0006 |
| Slain2 | 1,15 | 0,0006 |
| Aifm2 | 0,88 | 0,0006 |
| Tubb6 | 1,11 | 0,0006 |
| Ifi202 | 0,79 | 0,0006 |
| Ubqln1 | 1,19 | 0,0006 |
| Hyi | 2,38 | 0,0007 |
| Alyref | 1,76 | 0,0007 |
| Pts | 3,35 | 0,0007 |
| Fam98b | 1,11 | 0,0009 |
| Smug1 | 1,22 | 0,0009 |
| Npm1 | 1,18 | 0,0009 |
| Tmem33 | 0,82 | 0,0009 |
| Mrpl12 | 1,19 | 0,0009 |
| Med15 | 1,16 | 0,0009 |
| Gnb2 | 0,88 | 0,0009 |
| Rps15a | 0,81 | 0,0010 |
| Btf3l4 | 1,11 | 0,0010 |
| Rps16 | 0,84 | 0,0010 |
| Mppe1 | 1,23 | 0,0010 |
| Gnai3 | 0,84 | 0,0010 |
| Zdhhc17 | 1,12 | 0,0010 |
| Naa30 | 0,82 | 0,0011 |
| Pycr3 | 1,64 | 0,0012 |
| Tomm40 | 0,81 | 0,0012 |
| D11Wsu47e | 1,37 | 0,0012 |
| Rpl38 | 0,75 | 0,0012 |
| Acsl4 | 0,85 | 0,0012 |
| Anapc5 | 0,67 | 0,0013 |
| Czib | 1,16 | 0,0013 |
| Rps11 | 0,92 | 0,0013 |
| Smndc1 | 1,24 | 0,0013 |
| Smarca5 | 0,93 | 0,0013 |
| Snrpa1 | 1,15 | 0,0013 |
| Ap2m1 | 0,87 | 0,0013 |
| Lmna | 1,11 | 0,0014 |
| Ubqln2 | 1,25 | 0,0014 |
| Ilvbl | 0,88 | 0,0014 |
| Rps3 | 0,86 | 0,0015 |
| MACIR | 1,30 | 0,0015 |
| Rpia | 0,83 | 0,0015 |
| Naa10 | 0,90 | 0,0015 |
| Hsd17b12 | 0,80 | 0,0016 |
| Ifi35 | 1,12 | 0,0016 |
| Srp72 | 0,89 | 0,0016 |
| Anxa3 | 1,74 | 0,0017 |
| Gm28040;Golt1a | 0,49 | 0,0017 |
| Rpl7a | 0,88 | 0,0017 |
| Ppm1a | 1,13 | 0,0017 |
| Eif2b1 | 0,87 | 0,0018 |
| Rnf13 | 1,29 | 0,0019 |
| Slc31a1 | 0,66 | 0,0020 |
| Mydgf | 0,83 | 0,0020 |
| Rpl26 | 0,87 | 0,0020 |
| Rab35 | 0,87 | 0,0021 |
| Mta3 | 0,87 | 0,0021 |
| Rnf185 | 0,47 | 0,0021 |
| Wtap | 1,14 | 0,0022 |
| Abce1 | 0,94 | 0,0023 |
| Kdm3b | 1,07 | 0,0023 |
| Trappc2l | 0,85 | 0,0024 |
| Alg13 | 1,64 | 0,0024 |
| Rps24 | 0,82 | 0,0024 |
| Rpl30 | 0,78 | 0,0025 |
| Tpr | 1,08 | 0,0025 |
| Tfdp2 | 0,53 | 0,0025 |
| Elovl1 | 0,63 | 0,0025 |
| Tmem109 | 1,28 | 0,0026 |
| Eif3i | 0,92 | 0,0026 |
| Pdcd6 | 1,14 | 0,0026 |
| Mesd | 0,91 | 0,0026 |
| Eif4ebp1 | 1,28 | 0,0026 |
| Ddrgk1 | 1,25 | 0,0027 |
| Cnpy4 | 1,16 | 0,0027 |
| Lsm8 | 1,17 | 0,0027 |
| Snrpb | 0,80 | 0,0028 |
| Pigt | 0,86 | 0,0028 |
| Erh | 1,26 | 0,0028 |
| UPF0449 protein C19orf25 | 1,29 | 0,0028 |
| Myl12b | 1,11 | 0,0028 |
| Qrich1 | 1,15 | 0,0028 |
| Ppil3 | 1,14 | 0,0029 |
| Ppp1r12b | 1,30 | 0,0029 |
| Eif3f | 0,90 | 0,0030 |
| Copa | 0,91 | 0,0030 |
| Pyurf | 1,63 | 0,0030 |
| Cpped1 | 1,49 | 0,0030 |
| Rpl22 | 0,87 | 0,0031 |
| Cd63 | 3,47 | 0,0031 |
| Tox4 | 1,22 | 0,0031 |
| Ppp1r14b | 1,35 | 0,0032 |
| Rcsd1 | 1,18 | 0,0032 |
| Srsf1 | 1,13 | 0,0032 |
| Slc25a35 | 0,83 | 0,0033 |
| Prps1l3;Prps1 | 0,88 | 0,0033 |
| Ube2d2 | 1,19 | 0,0035 |

| | | |
|---|---|---|
| Rpl32 | 0,84 | 0,0035 |
| Mospd2 | 0,94 | 0,0037 |
| Polr2e | 0,88 | 0,0037 |
| Arfgef1 | 0,93 | 0,0038 |
| Rps18-ps6;Rps18 | 0,91 | 0,0038 |
| Dxo | 0,78 | 0,0038 |
| Ak2 | 1,09 | 0,0038 |
| Tmem53 | 0,68 | 0,0038 |
| Tor1a | 0,86 | 0,0039 |
| Elp2 | 0,89 | 0,0039 |
| Creld2 | 1,12 | 0,0039 |
| Hsd17b11 | 0,85 | 0,0039 |
| Oxa1l | 0,84 | 0,0039 |
| Erf | 0,72 | 0,0039 |
| Tlr9 | 0,90 | 0,0039 |
| Tmem256 | 1,40 | 0,0040 |
| Pdia3 | 1,08 | 0,0040 |
| Spcs3 | 0,83 | 0,0041 |
| Cpsf3 | 0,89 | 0,0041 |
| Prpf38b | 0,91 | 0,0042 |
| Mak16 | 0,84 | 0,0045 |
| Prkag1 | 0,82 | 0,0045 |
| Golgb1 | 1,08 | 0,0046 |
| Clec12a | 1,23 | 0,0046 |
| Psmd14 | 0,84 | 0,0046 |
| Gm29094 | 0,80 | 0,0047 |
| Drg2 | 0,85 | 0,0049 |
| Cmah | 1,39 | 0,0049 |
| Ldah | 0,87 | 0,0049 |
| Taf7 | 1,30 | 0,0049 |
| Rcbtb1 | 0,81 | 0,0049 |
| Ccdc97 | 1,26 | 0,0050 |
| Rhbdd2 | 0,71 | 0,0050 |
| Golga5 | 1,07 | 0,0051 |
| Tmppe | 0,84 | 0,0052 |
| Rps4x | 0,93 | 0,0052 |
| Tmem199 | 0,83 | 0,0052 |
| Ptpn9 | 0,90 | 0,0052 |
| Tmem260 | 2,03 | 0,0052 |
| Rpl21 | 0,88 | 0,0054 |
| Rad23b | 1,10 | 0,0054 |
| Pus7l | 1,23 | 0,0055 |
| Pigu | 0,77 | 0,0055 |
| Cux1 | 1,13 | 0,0055 |
| Tuba1c | 1,31 | 0,0055 |
| Cnn2 | 1,11 | 0,0055 |
| Atp1a4 | 1,75 | 0,0055 |
| Pam16 | 1,10 | 0,0056 |
| Med30 | 1,20 | 0,0056 |
| Stk26 | 1,19 | 0,0057 |
| Slc7a6os | 1,19 | 0,0057 |
| Lsm7 | 1,10 | 0,0058 |
| Fis1 | 0,81 | 0,0058 |
| Erlin2 | 0,92 | 0,0059 |
| Spcs2 | 0,83 | 0,0059 |
| Slc35c1 | 0,70 | 0,0060 |
| Zfp850 | 0,45 | 0,0061 |
| Bmp2k | 1,08 | 0,0061 |
| Tnrc6b | 1,17 | 0,0062 |
| Kcnn2 | 2,29 | 0,0063 |
| Asxl1 | 0,68 | 0,0063 |
| H3c1 | 0,85 | 0,0063 |
| Csde1 | 0,92 | 0,0064 |
| Sgsh | 2,74 | 0,0065 |
| Iws1 | 1,15 | 0,0065 |
| Alyref2 | 1,12 | 0,0065 |
| Wdr5 | 0,91 | 0,0065 |
| Hook3 | 1,13 | 0,0065 |
| Slc39a11 | 1,30 | 0,0065 |
| Tyw5 | 1,17 | 0,0066 |
| Numa1 | 1,08 | 0,0066 |
| Paics | 0,85 | 0,0067 |
| Lbh | 1,35 | 0,0070 |
| Ddx46 | 0,92 | 0,0071 |
| Snrnp200 | 0,93 | 0,0071 |
| Pcyox1 | 0,89 | 0,0071 |
| Aqp8 | 0,38 | 0,0071 |
| Ddt | 12,34 | 0,0072 |
| 0610009B22Rik | 0,80 | 0,0073 |
| Anp32b | 1,18 | 0,0075 |
| Madd | 1,67 | 0,0075 |
| Sel1l | 0,89 | 0,0076 |
| Rpl3 | 0,91 | 0,0076 |
| Exosc8 | 1,12 | 0,0076 |
| Tnip1 | 1,19 | 0,0078 |
| Gnaq | 0,92 | 0,0078 |
| Tap2 | 0,90 | 0,0078 |
| Fip1l1 | 1,17 | 0,0078 |
| Eif1b | 1,15 | 0,0078 |
| Atxn1 | 1,24 | 0,0079 |
| Ddx21 | 0,90 | 0,0079 |
| Ubqln4 | 1,15 | 0,0080 |
| Srd5a1 | 0,40 | 0,0080 |
| Oard1 | 0,82 | 0,0080 |
| Dnah7b;Dnah7a | 1,42 | 0,0081 |
| Hnf4a | 0,40 | 0,0081 |
| Dapk1 | 0,89 | 0,0082 |
| Hnrnpr | 1,31 | 0,0086 |
| Calm1;Calm2;Calm3 | 1,73 | 0,0086 |
| Cyp20a1 | 0,88 | 0,0087 |
| Wdr55 | 1,08 | 0,0087 |
| Bax | 1,18 | 0,0087 |
| N6amt1 | 1,16 | 0,0088 |
| Dpp8 | 0,91 | 0,0089 |
| Pnpt1 | 0,89 | 0,0089 |
| Tpd52l2 | 1,13 | 0,0090 |
| Ctdnep1 | 0,74 | 0,0090 |
| Dolpp1 | 1,29 | 0,0091 |
| Uxt | 1,25 | 0,0091 |
| Tbl1x | 0,79 | 0,0092 |
| Ubxn1 | 1,27 | 0,0095 |
| Dnajb11 | 1,07 | 0,0096 |
| Paat | 1,30 | 0,0096 |
| Tsfm | 1,11 | 0,0097 |
| Mettl21A | 0,74 | 0,0098 |
| Phf1 | 2,42 | 0,0101 |
| Bco2 | 0,27 | 0,0102 |
| Ubqln1 | 1,67 | 0,0102 |
| Mzb1 | 1,28 | 0,0102 |
| Gid8 | 1,15 | 0,0103 |
| Naa50 | 0,89 | 0,0103 |
| Anxa5 | 1,33 | 0,0103 |
| Caprin1 | 1,14 | 0,0104 |
| Fkbp4 | 0,93 | 0,0104 |
| Rbmx | 1,10 | 0,0104 |
| Rps8 | 0,90 | 0,0104 |
| Mrpl33 | 0,78 | 0,0106 |
| Tmbim6 | 1,41 | 0,0107 |
| Rab18 | 0,84 | 0,0109 |
| Tpr | 1,16 | 0,0110 |
| Tut1 | 1,13 | 0,0110 |
| Arl1 | 0,80 | 0,0111 |
| Anp32a | 1,10 | 0,0114 |
| Psmd3 | 0,90 | 0,0114 |
| Cdc42ep2 | 1,18 | 0,0114 |
| Tdp1 | 0,74 | 0,0115 |
| Sfr1 | 1,24 | 0,0115 |
| Nhlrc3 | 1,53 | 0,0117 |
| Sf3b5 | 1,19 | 0,0117 |
| Snd1 | 0,92 | 0,0117 |
| Slc25a19 | 0,84 | 0,0117 |
| Serinc3 | 0,73 | 0,0118 |
| Pygm | 0,73 | 0,0118 |
| Rpl23 | 0,88 | 0,0119 |
| Psmc6 | 1,08 | 0,0120 |
| Copb2 | 0,91 | 0,0121 |

| | | |
|---|---|---|
| Trappc6b | 0,85 | 0,0122 |
| Prkd1 | 7,45 | 0,0122 |
| Atxn7l3b | 1,25 | 0,0123 |
| Ncl | 1,10 | 0,0123 |
| Dcps | 1,09 | 0,0124 |
| Dyrk1a | 1,19 | 0,0125 |
| Vti1a | 1,07 | 0,0126 |
| Mrpl53 | 1,14 | 0,0127 |
| Pigg | 1,21 | 0,0127 |
| Alkbh3 | 1,12 | 0,0128 |
| Traf3ip2 | 1,69 | 0,0128 |
| Ubn2 | 1,25 | 0,0129 |
| Smarcb1 | 0,75 | 0,0129 |
| Olfr44 | 0,54 | 0,0130 |
| Aox1 | 1,31 | 0,0131 |
| Eif4h | 1,14 | 0,0131 |
| Smim9 | 3,32 | 0,0131 |
| Polr2b | 0,96 | 0,0131 |
| Ppp6c | 0,86 | 0,0132 |
| Thrap3 | 1,09 | 0,0132 |
| Cd59a | 1,82 | 0,0132 |
| Ogfr | 1,07 | 0,0132 |
| Tnfrsf14 | 1,43 | 0,0133 |
| Rdh13 | 0,87 | 0,0134 |
| Cbx1 | 1,18 | 0,0136 |
| Nsfl1c | 1,11 | 0,0138 |
| Calhm2 | 0,51 | 0,0138 |
| Hnrnpa2b1 | 1,06 | 0,0139 |
| Mettl2 | 1,22 | 0,0139 |
| Iqcf3 | 1,34 | 0,0142 |
| Chmp1b1 | 0,81 | 0,0142 |
| Ccdc124 | 1,25 | 0,0142 |
| Pet117 | 1,32 | 0,0142 |
| Numb | 1,47 | 0,0144 |
| Mrpl20 | 0,79 | 0,0145 |
| Cycs | 0,74 | 0,0146 |
| Morf4l1 | 0,91 | 0,0147 |
| Lpar6 | 1,50 | 0,0148 |
| Haus7 | 1,33 | 0,0148 |
| Fam98a | 0,85 | 0,0148 |
| Scfd1 | 1,05 | 0,0149 |
| Uchl3 | 1,11 | 0,0149 |
| Cyp4f16 | 0,68 | 0,0150 |
| Polr2g | 0,89 | 0,0151 |
| Rack1 | 0,90 | 0,0151 |
| Ube2h | 1,14 | 0,0152 |
| Sec24b | 1,69 | 0,0156 |
| Lims1 | 0,75 | 0,0156 |
| Glod4 | 1,12 | 0,0156 |
| Alg3 | 1,14 | 0,0156 |
| Mtrex | 0,93 | 0,0157 |
| Sun3 | 1,97 | 0,0157 |
| Aatf | 1,15 | 0,0157 |
| Rps28 | 1,21 | 0,0158 |
| Acta1 | 1,35 | 0,0158 |
| Fyttd1 | 1,12 | 0,0160 |
| Fau | 0,82 | 0,0162 |
| Ranbp10 | 1,10 | 0,0162 |
| Dhrs7b | 0,83 | 0,0163 |
| Serinc1 | 0,78 | 0,0164 |
| Vamp4 | 1,20 | 0,0164 |
| Tmed7 | 0,82 | 0,0166 |
| Sf3b3 | 0,90 | 0,0166 |
| Safb2 | 1,10 | 0,0168 |
| Cdkn2aip | 1,09 | 0,0169 |
| AF366264 | 2,78 | 0,0170 |
| Tmed9 | 1,12 | 0,0170 |
| Ppib | 0,88 | 0,0171 |
| Clp1 | 0,87 | 0,0172 |
| Git2 | 1,64 | 0,0173 |
| Rab24 | 0,91 | 0,0174 |
| Ifi204 | 0,87 | 0,0175 |
| Phf21a | 0,77 | 0,0175 |
| Med25 | 0,87 | 0,0177 |
| Chchd1 | 1,27 | 0,0177 |
| Ccdc90b | 1,10 | 0,0178 |
| Mettl3 | 1,20 | 0,0178 |
| Sarnp | 1,10 | 0,0179 |
| Slc48a1 | 0,58 | 0,0179 |
| Abcc1 | 0,85 | 0,0181 |
| Lamtor2 | 0,82 | 0,0181 |
| Sgms1 | 1,21 | 0,0183 |
| Card10 | 0,74 | 0,0184 |
| Zbp1 | 1,88 | 0,0184 |
| Nme3 | 0,75 | 0,0184 |
| Shkbp1 | 2,30 | 0,0185 |
| Snrpd1 | 0,82 | 0,0185 |
| M6pr | 1,11 | 0,0186 |
| Ngrn | 1,19 | 0,0186 |
| Rab22a | 0,87 | 0,0186 |
| Hmgcr | 0,70 | 0,0187 |
| Tenm4 | 0,67 | 0,0187 |
| Snrpe | 0,87 | 0,0188 |
| Rragc | 0,75 | 0,0188 |
| Camkk1 | 0,89 | 0,0188 |
| Idh3g | 0,86 | 0,0189 |
| Psma3 | 0,85 | 0,0189 |
| Map2k2 | 0,91 | 0,0190 |
| Kpna1 | 0,93 | 0,0190 |
| Cdk13 | 1,14 | 0,0190 |
| Cep63 | 1,41 | 0,0191 |
| Atrn | 0,81 | 0,0191 |
| Tmem167;Tmem167a | 0,84 | 0,0192 |
| Clk4 | 0,79 | 0,0192 |
| Mob3a | 0,75 | 0,0193 |
| Ppp2cb | 1,24 | 0,0196 |
| Cwc27 | 1,10 | 0,0196 |
| Mrpl21 | 0,90 | 0,0197 |
| Fth1 | 3,37 | 0,0199 |
| Sf3a3 | 1,10 | 0,0199 |
| Eps15 | 1,12 | 0,0200 |
| Rab20 | 0,87 | 0,0200 |
| Srp9 | 0,82 | 0,0200 |
| Stambpl1 | 0,86 | 0,0201 |
| Cdc37l1 | 1,15 | 0,0203 |
| Lage3 | 0,88 | 0,0204 |
| Magoh | 0,81 | 0,0204 |
| Cnpy3 | 1,13 | 0,0204 |
| Twf2 | 0,82 | 0,0205 |
| Prkaa1 | 0,91 | 0,0206 |
| Slc38a10 | 1,35 | 0,0207 |
| Rbm14 | 1,10 | 0,0207 |
| Casp4 | 0,84 | 0,0207 |
| Mfsd5 | 0,81 | 0,0208 |
| Tmpo | 1,11 | 0,0209 |
| Ak2 | 1,24 | 0,0209 |
| Mau2 | 0,91 | 0,0209 |
| Eef1g | 0,94 | 0,0211 |
| Pfdn4 | 1,15 | 0,0212 |
| Fyco1 | 1,10 | 0,0213 |
| Fnbp1l | 1,17 | 0,0213 |
| Ccdc28a | 0,73 | 0,0214 |
| Polr2d | 1,23 | 0,0215 |
| Timp2 | 2,31 | 0,0216 |
| Hnrnpk | 1,07 | 0,0217 |
| Tecpr2 | 0,87 | 0,0218 |
| Rab3d | 0,86 | 0,0220 |
| Eed | 0,88 | 0,0220 |
| Ppp6r3 | 1,09 | 0,0223 |
| Pdia4 | 1,06 | 0,0223 |
| Txndc12 | 1,07 | 0,0223 |
| Ids | 1,14 | 0,0223 |
| Ilf2 | 0,92 | 0,0224 |
| Pfkfb3 | 0,85 | 0,0225 |
| Mtap | 0,92 | 0,0225 |
| Prpsap1 | 0,92 | 0,0227 |

| | | |
|---|---|---|
| Kcnab1 | 0,62 | 0,0229 |
| Slc66a3 | 0,71 | 0,0230 |
| Tmed8 | 0,76 | 0,0230 |
| Fubp3 | 1,07 | 0,0231 |
| Exosc6 | 1,12 | 0,0231 |
| Dcxr | 0,82 | 0,0232 |
| Stx16 | 1,15 | 0,0233 |
| Ankrd40 | 1,26 | 0,0233 |
| Ascc1 | 0,85 | 0,0234 |
| Wac | 1,11 | 0,0234 |
| Rnf146 | 1,50 | 0,0234 |
| Tlcd1 | 0,73 | 0,0234 |
| Nup205 | 0,94 | 0,0238 |
| Stx6 | 1,18 | 0,0238 |
| Rras | 0,89 | 0,0238 |
| Gpd2 | 0,89 | 0,0240 |
| Cbfb | 1,12 | 0,0240 |
| Gmeb2 | 1,22 | 0,0242 |
| Rplp0 | 0,91 | 0,0242 |
| Kif2a | 1,08 | 0,0244 |
| Gpr89 | 0,82 | 0,0244 |
| Tnfsfm13;Tnfsf13 | 1,54 | 0,0245 |
| Pym1 | 1,44 | 0,0246 |
| Ftl1 | 3,68 | 0,0249 |
| Ythdc1 | 0,73 | 0,0249 |
| Hdgf | 1,11 | 0,0250 |
| Mrpl27 | 1,21 | 0,0250 |
| Prkab1 | 0,96 | 0,0252 |
| Bak1 | 0,88 | 0,0253 |
| Dpy19l4 | 1,31 | 0,0253 |
| Mrpl49 | 1,14 | 0,0255 |
| Pign | 1,15 | 0,0256 |
| Fam32a | 1,19 | 0,0256 |
| Ppp4r3a | 0,92 | 0,0257 |
| Noc3l | 1,14 | 0,0257 |
| Skap2 | 1,11 | 0,0258 |
| Lrrfip1 | 1,24 | 0,0258 |
| Jak3 | 1,18 | 0,0259 |
| Ftl1 | 3,52 | 0,0259 |
| Rbx1 | 0,83 | 0,0261 |
| Dis3 | 0,91 | 0,0262 |
| Vps8 | 0,93 | 0,0262 |
| Mnat1 | 1,12 | 0,0262 |
| Ireb2 | 1,17 | 0,0263 |
| Itch | 0,91 | 0,0264 |
| Hmgn2 | 0,35 | 0,0264 |
| Ighmbp2 | 0,82 | 0,0265 |
| Nectin2 | 1,10 | 0,0267 |
| Nap1l4 | 1,13 | 0,0267 |
| Avl9 | 0,88 | 0,0268 |
| Cds1 | 0,74 | 0,0268 |
| Rai14 | 1,06 | 0,0268 |
| Derl2 | 0,78 | 0,0269 |
| Prmt1 | 0,84 | 0,0270 |
| Araf | 0,89 | 0,0270 |
| Nif3l1 | 0,90 | 0,0272 |
| Pbp2 | 1,24 | 0,0272 |
| Manf | 1,08 | 0,0273 |
| Gstp2 | 0,25 | 0,0276 |
| Clcn2 | 0,69 | 0,0277 |
| Myo10 | 0,85 | 0,0278 |
| Calr | 1,09 | 0,0280 |
| Slc25a5 | 0,88 | 0,0280 |
| Plekho1 | 1,16 | 0,0280 |
| Acp3 | 1,39 | 0,0280 |
| Eef1a1 | 0,92 | 0,0280 |
| Dmac1 | 0,78 | 0,0282 |
| Rbm6 | 1,12 | 0,0284 |
| Snx16 | 1,25 | 0,0286 |
| Cyhr1 | 0,88 | 0,0289 |
| Atp13a1 | 0,95 | 0,0291 |
| Stk38 | 0,83 | 0,0291 |
| Nudt3 | 0,82 | 0,0291 |
| Mb21d2 | 0,69 | 0,0291 |
| Cfdp1 | 1,12 | 0,0292 |
| Rragd | 0,81 | 0,0293 |
| Prkci | 0,79 | 0,0293 |
| Pdcl3 | 1,18 | 0,0293 |
| Rab2b | 0,86 | 0,0295 |
| Kdm2a | 0,87 | 0,0296 |
| Rel | 0,87 | 0,0297 |
| Saal1 | 1,14 | 0,0297 |
| Bola1 | 1,14 | 0,0297 |
| Cdc42 | 1,14 | 0,0300 |
| Ywhaq | 1,14 | 0,0301 |
| Ubr2 | 0,91 | 0,0302 |
| Ints11 | 0,82 | 0,0302 |
| Mto1 | 0,76 | 0,0304 |
| Tspo | 0,77 | 0,0304 |
| Supt7l | 0,78 | 0,0305 |
| Snrpf | 0,86 | 0,0305 |
| Cops6 | 0,86 | 0,0309 |
| Sash1 | 1,08 | 0,0311 |
| Fbxl22 | 0,81 | 0,0312 |
| Rpl8 | 0,91 | 0,0314 |
| Dsc2 | 0,62 | 0,0316 |
| Dcp2 | 0,84 | 0,0317 |
| Mocs3 | 1,09 | 0,0319 |
| Skap1 | 1,41 | 0,0322 |
| Paxx | 1,17 | 0,0323 |
| Dip2a | 0,86 | 0,0323 |
| Ghdc | 0,89 | 0,0323 |
| Sult2a1 | 0,40 | 0,0324 |
| Agtpbp1 | 0,58 | 0,0324 |
| Dpy30 | 1,15 | 0,0324 |
| Dhrs7 | 0,88 | 0,0325 |
| Tango2 | 1,12 | 0,0326 |
| Arfgap2 | 1,07 | 0,0326 |
| Pcnp | 1,11 | 0,0327 |
| Mpeg1 | 1,23 | 0,0329 |
| Fgfr1op2 | 1,20 | 0,0329 |
| Mrpl14 | 0,84 | 0,0330 |
| Polr1f | 1,13 | 0,0330 |
| Plekhf1 | 0,74 | 0,0333 |
| Tmem63a | 0,88 | 0,0333 |
| Plac8 | 1,52 | 0,0334 |
| Trex1 | 1,08 | 0,0336 |
| Tmcc3 | 1,45 | 0,0336 |
| Rbm5 | 1,08 | 0,0337 |
| Psmg2 | 0,91 | 0,0339 |
| Sdhaf4 | 1,20 | 0,0339 |
| Ca2 | 0,86 | 0,0342 |
| Ppp2r1b | 1,09 | 0,0343 |
| Tlr2 | 0,79 | 0,0345 |
| Tmem120a | 0,86 | 0,0345 |
| Wipi2 | 0,89 | 0,0346 |
| Fus | 1,13 | 0,0347 |
| Ap2a2 | 0,93 | 0,0350 |
| Ddx24 | 1,08 | 0,0350 |
| Hdac10 | 0,76 | 0,0353 |
| Lamc1 | 1,84 | 0,0353 |
| Gm17728 | 1,52 | 0,0356 |
| Rbm8a | 1,14 | 0,0359 |
| Dek | 1,12 | 0,0359 |
| Exosc10 | 0,94 | 0,0361 |
| Ncdn | 0,90 | 0,0363 |
| Rbpj | 0,82 | 0,0365 |
| Ifngr1 | 1,26 | 0,0366 |
| Gemin5 | 0,90 | 0,0366 |
| Mrpl50 | 1,11 | 0,0367 |
| Sar1a | 0,87 | 0,0368 |
| Trmt2a | 1,07 | 0,0368 |
| Otud6b | 1,16 | 0,0369 |
| Myh9 | 1,07 | 0,0372 |
| Hdac2 | 0,92 | 0,0373 |
| Sart1 | 1,10 | 0,0374 |

| | | |
|---|---|---|
| Spcs1 | 0,89 | 0,0374 |
| Nat10 | 0,92 | 0,0374 |
| Gp49a | 0,72 | 0,0375 |
| Ilkap | 1,12 | 0,0375 |
| Emc1 | 0,88 | 0,0376 |
| Aptx | 0,74 | 0,0376 |
| Tmed1 | 1,17 | 0,0377 |
| Smad2 | 1,09 | 0,0378 |
| Scd2 | 0,56 | 0,0380 |
| Dguok | 0,85 | 0,0380 |
| Fkbp3 | 1,10 | 0,0381 |
| Map4k4 | 0,90 | 0,0381 |
| Erbin | 1,08 | 0,0386 |
| Bcl2l11 | 1,22 | 0,0386 |
| Tprg1l | 0,92 | 0,0387 |
| Parp10 | 0,93 | 0,0387 |
| Pomp | 1,19 | 0,0387 |
| Luc7l2 | 1,11 | 0,0388 |
| Col4a2 | 1,25 | 0,0388 |
| Acte1 | 2,24 | 0,0388 |
| Pdlim2 | 1,13 | 0,0388 |
| Sh3bgrl3 | 1,24 | 0,0391 |
| Emp3 | 1,29 | 0,0391 |
| Yipf5 | 1,35 | 0,0391 |
| Son | 1,05 | 0,0392 |
| Asrgl1 | 1,14 | 0,0393 |
| Limd1 | 1,18 | 0,0394 |
| Naa38 | 1,18 | 0,0394 |
| Papola | 0,89 | 0,0395 |
| Mdc1 | 0,86 | 0,0395 |
| Med1 | 0,89 | 0,0399 |
| Sec61a2 | 0,77 | 0,0400 |
| Rab11fip1 | 1,13 | 0,0401 |
| Alg5 | 0,92 | 0,0402 |
| Atg13 | 1,39 | 0,0403 |
| Mstn | 1,44 | 0,0403 |
| Acot3 | 0,42 | 0,0404 |
| Cwf19l2 | 0,67 | 0,0404 |
| Klhl36 | 0,69 | 0,0404 |
| Tgfbrap1 | 0,88 | 0,0404 |
| Srsf9 | 1,08 | 0,0404 |
| Slc25a3 | 0,79 | 0,0406 |
| Cd99l2 | 1,28 | 0,0408 |
| Taf10 | 1,18 | 0,0408 |
| Stip1 | 1,07 | 0,0409 |
| Cand2 | 1,69 | 0,0409 |
| Spata5l1 | 0,89 | 0,0409 |
| Tap1 | 0,91 | 0,0409 |
| Isyna1 | 1,38 | 0,0412 |
| Cpsf6 | 1,07 | 0,0412 |
| Myg1 | 1,06 | 0,0413 |
| Cmklr1 | 0,72 | 0,0414 |
| Wdr75 | 0,83 | 0,0414 |
| Ankrd27 | 1,36 | 0,0417 |
| Rps7 | 0,86 | 0,0417 |
| Tmem175 | 1,41 | 0,0420 |
| Ddx10 | 0,87 | 0,0420 |
| Nudt2 | 0,90 | 0,0421 |
| Ncbp3 | 1,12 | 0,0422 |
| Acin1 | 1,08 | 0,0424 |
| Itpkc | 1,33 | 0,0425 |
| Odr4 | 0,89 | 0,0425 |
| Pelo | 0,86 | 0,0426 |
| Zc3h15 | 1,08 | 0,0427 |
| Ddi2 | 1,13 | 0,0431 |
| Ogt | 1,23 | 0,0431 |
| A630001G21Rik | 1,22 | 0,0432 |
| Met | 0,83 | 0,0432 |
| Elp1 | 0,95 | 0,0433 |
| Phf8 | 1,17 | 0,0433 |
| Set | 0,40 | 0,0434 |
| Snap29 | 1,21 | 0,0436 |
| Tmem147 | 0,73 | 0,0436 |
| Psma6 | 0,87 | 0,0439 |
| Tfcp2 | 1,18 | 0,0445 |
| Zgpat | 0,81 | 0,0445 |
| Mphosph10 | 1,11 | 0,0446 |
| Hnrnpab | 1,12 | 0,0447 |
| H2-Aa | 1,33 | 0,0450 |
| Stk16 | 1,16 | 0,0450 |
| Epn2 | 1,12 | 0,0451 |
| Tmsb10 | 1,24 | 0,0451 |
| Scamp3 | 1,42 | 0,0453 |
| Crp | 0,66 | 0,0453 |
| Rnase4 | 0,54 | 0,0460 |
| Aplp2 | 1,25 | 0,0462 |
| Ms4a4b | 0,68 | 0,0466 |
| Eif4a2 | 0,92 | 0,0466 |
| Ipo7 | 0,90 | 0,0467 |
| Tmem63b | 0,84 | 0,0467 |
| Rbm12 | 1,08 | 0,0470 |
| Vps26b | 0,90 | 0,0470 |
| Fiz1 | 1,43 | 0,0471 |
| Bscl2 | 1,20 | 0,0471 |
| Tcerg1 | 1,06 | 0,0472 |
| Kansl1 | 0,68 | 0,0473 |
| Ccdc43 | 1,16 | 0,0475 |
| Pogz | 0,82 | 0,0477 |
| Slc25a43 | 1,37 | 0,0477 |
| Acbd6 | 1,19 | 0,0477 |
| Eml4 | 0,93 | 0,0478 |
| Mocos | 0,96 | 0,0481 |
| Scarf2 | 0,57 | 0,0483 |
| 9930111J21Rik1 | 1,11 | 0,0483 |
| Rptor | 0,92 | 0,0483 |
| Sltm | 1,31 | 0,0484 |
| Fbxo38 | 1,13 | 0,0484 |
| Mon2 | 0,95 | 0,0485 |
| Parp12 | 0,82 | 0,0487 |
| Klhl25 | 0,88 | 0,0488 |
| Nfatc3 | 0,70 | 0,0488 |
| Lztfl1 | 1,16 | 0,0488 |
| Narf | 0,66 | 0,0490 |
| Dnm2 | 0,81 | 0,0492 |
| Mapre3 | 1,39 | 0,0492 |
| Sin3b | 0,71 | 0,0493 |
| Usp46 | 0,83 | 0,0493 |
| Rheb | 0,92 | 0,0493 |
| Cfp | 1,13 | 0,0494 |
| Rpl31 | 0,91 | 0,0496 |
| Prep | 0,89 | 0,0497 |
| Slc2a3 | 0,82 | 0,0498 |
| Metap2 | 1,33 | 0,0499 |
| Slc3a1 | 0,82 | 0,0499 |
| Aig1 | 0,75 | 0,0499 |
